# Actinium-225 dendrimer-radioconjugates combined with low-dose standard-of-care chemotherapy: site-independent treatment of triple negative breast cancer metastases

**DOI:** 10.1101/2025.08.25.670941

**Authors:** Pooja Hariharan, Rajiv Nair, Aira Sarkar, Kathleen L. Gabrielson, Tony Wu, Chang Liu, Wathsala Liyanage, Rangaramanujam M Kannan, Stavroula Sofou

**Affiliations:** Chemical and Biomolecular Engineering (ChemBE), Institute for NanoBioTechnology (INBT), Johns Hopkins University, Baltimore, MD, USA; Molecular and Comparative Pathobiology, Johns Hopkins University, Baltimore, MD, USA; Center for Nanomedicine at the Wilmer Eye Institute, Johns Hopkins University, Baltimore, MD, USA; Sidney Kimmel Comprehensive Cancer Center, Cancer Invasion & Metastasis Program, Department of Oncology, Johns Hopkins University, Baltimore, MD, USA

**Keywords:** actinium-225, alpha-particle radionuclide therapy, dendrimer-radioconjugates, triple negative breast cancer, spheroids, cisplatin

## Abstract

**PURPOSE:** Metastatic triple negative breast cancer (mTNBC) is incurable largely due to the development of drug resistance, the lack of selective cell targeting, and/or limitations in tumor drug delivery that vary depending on the different (metastatic) tumor locations.

**METHODS:** The potential of a single type of systemic, targeted alpha-particle therapy (TAT) was investigated for addressing the above challenges of TNBC tumors implanted at different anatomic sites in mice. Actinium-225 dendrimer-radioconjugates alone, and/or after pretreatment with low-dose standard-of-care cisplatin, were assessed *in vitro* and on immune-competent 4T1-Balb/c mouse models with tumors implanted intracranially, orthotopically or subcutaneously.

**RESULTS:** *In vitro*, TAT’s efficacy was *enhanced by cisplatin*. *In vivo*, treatment was initiated well after tumors had grown (V_to_ = 39 ± 14 mm^3^ in the intracranial model, and V_to_=100mm^3^ in the orthotopic and subcutaneous models). Across all tumor implantation sites, a unified correlation was observed between animal mean survival and the dendrimer-delivered tumor absorbed doses, which were selectively *increased by low-dose cisplatin pretreatment*. Importantly, in all animal models, the mean survival following systemic treatment with both modalities was *significantly longer* vs. each modality alone and/or vs. no treatment, at injected doses that did not cause long-term (10-month) toxicities in tumor-free mice.

**CONCLUSION:** Systemically-injected dendrimer-delivered TAT, *combined with low-dose cisplatin pretreatment*, can safely extend survival independent of mTNBC tumors’ anatomic site, potentially presenting a single type of therapy to simultaneously treat multi-site mTNBC.

## Introduction

Almost forty-five percent of patients diagnosed with triple negative breast cancer (TNBC) will ultimately progress to metastatic disease [1]. The most common soft-tissue sites of TNBC metastases include the brain, liver and lung [2]. Advanced, metastatic TNBC (mTNBC) is difficult to treat effectively largely due to the development of resistance to approved agents and the lack of (conventional) molecular targets being overexpressed by and/or being unique to TNBC cells, that limit the ability to selectively deliver lethal doses of therapeutic agents to tumors [1]. Furthermore, in some TNBC metastatic sites, such as the brain, there are additional limitations in effective tumor delivery of systemically injected treatments [3]. Consequently, current treatments extend survival by only a few months [1, 2]. Therefore, development of treatments for mTNBC, effectively acting on different metastatic tumor sites, is urgently needed.

Targeted alpha-particle therapies (TAT) have been impressively successful in “targetable”, difficult-to kill cancers [4] and are currently being evaluated in clinical trials against a variety of soft-tissue solid tumors [5]. The enthusiasm for TAT is due to the unparalleled killing efficacy of, and irradiation precision by, alpha-particles (α-particles) enabled by the selectivity of tumor targeting technologies. Due to their high energy (MeV), α-particles cause complex double-strand DNA breaks as they traverse the cell nucleus. The inability to repair this DNA damage is the reason why α-particles [6] are generally impervious to resistance [7, 8], independent of cell origin and/or resistance to other agents. Notably, α-particles travel only a few cell lengths in tissue (40-80 μm), therefore minimizing irradiation of the healthy tissue(s) surrounding the targeted tumors.

To deliver TAT to mTNBC model tumors implanted at different anatomic sites, we investigated nanometer-sized dendrimers, which are highly branched polymers with precisely adjustable size [9]. We employed the hydroxyl-terminated generation-6 dendrimers (of ∼7 nm in diameter) that were previously reported to extensively associate with cancer cells, brain tumors and tumor-associated macrophages [10, 11], while sparing resting macrophages and healthy brain [11]. Although the vascular permeability of established, soft-tissue solid tumors to nanotherapeutics may not always present a major limitation in delivery, this usually becomes a major challenge with systemic therapies aiming to deliver treatments to tumors in the brain [3]. Importantly, systemically-injected hydroxyl PAMAM dendrimers have been demonstrated to cross the blood brain tumor barrier in mouse and rat brain-tumor models [10, 11], and dendrimer-drug conjugates have shown promise in multiple Phase 1 and Phase 2 trials, both on safety and efficacy measures, including addressing neurological consequences of severe COVID-19 [12, 13]. Therefore, building on the promise of hydroxyl PAMAM dendrimers ability to selectively target cancer cells, tumors in the brain and TAM from systemic delivery, we investigated their potential of becoming a *general vector* of TAT against established, soft-tissue TNBC metastases.

In this proof-of-concept study, generation 6 hydroxyl PAMAM dendrimers were radiolabeled with the α-particle emitter actinium-225 and were demonstrated to selectively associate with murine 4T1 TNBC cells *in vitro*, on monolayers and spheroids. Additionally, we investigated the ability of systemically-injected dendrimer-radioconjugates to selectively deliver lethal TAT doses to syngeneic 4T1 TNBC tumors implanted in the brain, subcutaneously or in the mammary fat pad of Balb/c mice. These sites were chosen to represent different metastatic and/or recurrent tumor sites. We evaluated the long-term toxicities of dendrimer-radioconjugates on tumor-free mice, the radiation dosimetry in each of the tumor-bearing mouse models, the dendrimer-radioconjugates’ therapeutic advantage in the presence and absence of pretreatment with low-dose cisplatin - employed as a standard of care for TNBC - and we also investigated potential correlations between animal survival and tumor absorbed doses for different treatment modalities and tumor implantation sites.

## Materials and Methods

Materials’ and reagents’ sourcing is detailed in the Supporting Information. The actinium-225 used in this research was supplied by the U.S. Department of Energy Isotope Program managed by the Office of Isotope R&D and Production.

### Cell lines

The cell line 4T1 was a gift from Dr. Daniele Gilkes at Johns Hopkins University, and was cultured in Roswell Park Memorial Institute (RPMI) media, supplemented with 10% FBS, 100 units/mL Penicillin and 100 µg/mL Streptomycin in an incubator at 37 °C and 5% CO_2_

### Dendrimer labeling and characterization

Hydroxyl-terminated Generation-6 poly(amidoamine) dendrimers were conjugated with the chelator DOTA (DOTA-dendrimer) or the chelator DTPA (DTPA-dendrimer) or the fluorescent label Cyanine 5 (Cy5-dendrimers), as preciously described [10, 11]. Size and zeta potential were measured by a Nanoseries Zetasizer (Malvern Instruments Ltd, Worcestershire, UK).

To radiolabel DOTA-dendrimers with [^225^Ac]Ac or DTPA-dendrimers with [^111^In]In, a one-step radiolabeling protocol was followed, and the specific activity and stability of radiolabeling were evaluated as previously described [11, 14]. The stability of radiolabeling was assessed 24 hours after incubating dendrimer-radioconjugates in serum-supplemented media at pH 7.4 and 37°C.

### Clonogenic Survival Assay

Five hundred thousand 4T1 cells were plated per well in a 6-well plate and were treated for 6 hours with each modality alone or together. Following treatment and washing, cells were seeded in tissue culture dishes to allow for colony formation, as previously described [11, 14]. The survival fraction was evaluated approximately after 12 cell-doubling times (doubling-time = 14 hours) and was calculated by dividing the number of colonies for each treatment group to the non-treated control group accounting for plating efficiency.

To calculate the mean activity associated per cell after completion of incubation, at the higher activity concentrations, the activity (γ-photon emissions of bismuth-213) associated with a fraction of cells, pooled from the parent suspension, was divided by the number of live cells in the same fraction, determined by Trypan Blue.

### Dendrimer association with 4T1 cells

A BD FACSCanto Flow cytometer (BD Biosciences, San Jose, CA, USA) was employed to analyze the extent of Cy5-dendrimers’ association with 4T1 cells following their incubation on ice for 1 hr (10×10⁶ cells/mL incubated at a dendrimer-to-cell ratio of 50×10⁶:1). Additionally, to evaluate the potential effect of chemotherapy on the kinetics and extent of dendrimer cell association, cells (2 million cells/mL) were incubated with Cy5-dendrimer (20.4 g/mL, and dendrimer-to-cell ratio of 50×10⁶:1) at 37 °C and 4 °C with or without chemotherapy, and at different time points 1 mL of the parent suspension was removed, centrifuged, and washed thrice with ice cold PBS before the cell associated fluorescence was measured using a Fluorolog-3 Fluorometer (Horiba Instruments, Edison, NJ).

### Spheroid Studies

Spheroids were formed by plating 3,000 4T1 cells per well onto ultra-low adhesion U-shaped 96-well plates following centrifugation at 1,023 RCF for 10 minutes and were then let grow until they reached 400μm in diameter.

To measure the interstitial pH gradient, spheroids were incubated for 12 hours (overnight) with 200 μM SNARF-4F, a membrane-impermeant pH-sensitive dye (excitation:488nm; emission: 580 nm and 640 nm). The ratios of fluorescence intensities at the two emission wavelengths vary with pH [15], and upon calibration, were employed to calculate the interstitial pH as a function of spheroid radius. After incubation, spheroids were washed with fresh media and imaged using an Olympus FV4000 Confocal Microscope at 10x zoom (Evident Scientific, Tokyo, Japan). On the images of the spheroids’ equatorial optical slices, an in-house erosion algorithm was applied to obtain the radial average intensities for both emission channels, and then their ratios were computed. These ratios were converted into radial pH values using a calibration curve obtained by imaging 50 μM SNARF-4F in media of varying pH in the range of interest (6.0–7.4).

To evaluate the dendrimers’ and cisplatin’s spatiotemporal microdistributions in spheroids, the latter were incubated with 10 μg/mL Cy5-dendrimers (ex/em: 651/670 nm) or 10μM CFDA-SE (ex/em: 492/517 nm) – used as a fluorescent surrogate of cisplatin - for up to 3 hours or 15 minutes, respectively. At different time points during the “uptake phase” (during incubation) and “clearance phase” (post incubation, when spheroids were transferred to fresh media), different spheroids were fished, embedded in Cryochrome™ gel and snap-frozen over dry ice before cryosectioning 20 μm thick equatorial sections, using a HM550 cryotome (Thermo Scientific, Waltham, MA) that were imaged using a Zeiss LSM-780 Laser Scanning Confocal Microscope (Zeiss, White Plains, NJ). As above, the radial microdistributions were analyzed using an erosion code. Fluorescence intensities were converted to concentrations using a calibration curve generated from serial dilutions of the fluorophores in a 20 μm-pathlength cuvette, imaged with the same microscope. Spheroids not incubated with fluorophores were used to establish background fluorescence levels.

To assess the effect of treatments on spheroid regrowth, post incubation of spheroids with [²²⁵Ac]Ac-DOTA-dendrimers and/or cisplatin, spheroids were transferred to an ultra-low attachment U-bottom plate containing fresh media. Their size was monitored daily for approximately 4 days. At that point, each spheroid was transferred to a separate well of a 48-well flat-bottom adherent plate and allowed to grow until cells from the non-treated group reached 70–80% confluency. At that point cells from all groups were then trypsinized, stained with trypan blue, and counted using a hemocytometer. Cell counts were normalized to the non-treated condition to calculate the “percent spheroid regrowth relative to no treatment.”

### Animal studies

Mice were housed in filter top cages with sterile food and water, and all studies were performed in compliance with the Institutional Animal Care and Use Committee protocol guidelines. Four- to six-week-old, 18g BALB/c female mice (Jackson Laboratory, Bar Harbor, ME, USA), were inoculated with 500 4T1 cells in 2 µL Matrigel^TM^ (Corning, NY, USA) either intracranially as previously described [10, 11], or with 10,000 4T1 cells in 100µL of Matrigel^TM^ in the mammary fat pad or subcutaneously.

Biodistributions were evaluated by intravenously administering 100µL of 740kBq [^111^In]In-DTPA-dendrimers. At various time points, mice were sacrificed, and tumors and normal-organs were excised, weighted, and a γ-counter was used to measure their associated activity, which was then decay-corrected and reported as percentage of the initially injected activity per gram of tissue (%IA/g).

Treatments were administered on day 10 after tumor inoculation (intracranial model) or when the mammary or the subcutaneous tumor volume reached 100mm^3^. When indicated, cisplatin (100µL of 5mg CDDP/kg mouse) was administered intraperitoneally 24 hours before intravenous injection of [^225^Ac]Ac-DOTA-dendrimers in 100µL. Following treatment, weights of mice were measured, and tumor volume (in the mammary and subcutaneous models) was calculated using the ellipsoid formula V=4*π*α*β^2^/3, where α and β are the major and minor diameters, respectively, measured by a digital caliper. Animals were euthanized when weight loss reached ≥20%, if tumor size was greater than 10% of body weight at the day of injection of therapy, or if the tumor interfered with the movement or function of mice. Following euthanasia, tumors and normal organs were harvested, fixed, stained with H&E, and evaluated for pathological findings.

### Dosimetry

The biodistributions of [^111^In]In-DTPA-dendrimers were utilized to calculate the absorbed doses for [^225^Ac]Ac-DOTA-dendrimers, assuming that all radioactive daughters were retained at the site of the parent decay, as previously described (*8*), using the software package RAPID Dosimetry (Baltimore, MD).

### Statistical Analysis

Results were reported as the arithmetic mean of n independent measurements ± the standard deviation. One-way ANOVA and/or the unpaired Student’s t test were used to calculate differences between experimental groups. Comparison of survival on Kaplan-Meier plots was performed using the log-rank test. P-values <0.05 were considered significant. * indicates 0.01<p-values<0.05; **<0.01, ***<0.001.

## Results

### Characterization of dendrimer-radioconjugates

Well-defined dendrimer-DOTA conjugates to enable actinium chelation were prepared and characterized using a previously described protocol [11]. The dendrimer-conjugates were radiolabeled with actinium-225 ([^225^Ac]Ac-DOTA-dendrimer) or indium-111 ([^111^In]In-DTPA-dendrimer) through chelation and were stable, with high radiochemical purity and reasonable specific activity (Table 1). Zeta potential measurements indicated a near-neutral surface charge for [^225^Ac]Ac-DOTA-dendrimer, with a value of - 0.16 ± 0.12 mV (n=3).

**Table 1.** Characterization of dendrimer-radioconjugates employed throughout this study. * Added activity for actinium-225 chelation was 5550kBq (150μCi). ** Added activity for indium-111 chelation was 148MBq (4mCi). Shown are the mean values ± the standard deviation of n independent measurements. Stability was assessed as % retention of activity by the radioconjugates following a 24-hour incubation in media at 37°C.

| $[^{225}\text{Ac}]\text{Ac-DOTA-dendrimer}$ | % Radiolabeling Efficiency | Specific Activity (kBq/mg dendrimer)* | Specific Activity (kBq/nmol dendrimer) | % Radiochemical Purity | % Stability |
| --- | --- | --- | --- | --- | --- |
| n=11 | $24.7 \pm 2.2$ | $1373 \pm 120$ | $80 \pm 7$ | $94.4 \pm 0.9$ | $92.5 \pm 1.1$ |
| $[^{111}\text{In}]\text{In-DTPA-dendrimer}$ | % Radiolabeling Efficiency | Specific Activity (kBq/mg dendrimer)** | Specific Activity (kBq/nmol dendrimer) | % Radiochemical Purity | % Stability |
| n=2 | $51.9 \pm 3.1$ | $105209 \pm 7247$ | $6134.6 \pm 422.6$ | $93.1 \pm 0.4$ | $93.9 \pm 1.1$ |

### Low-dose standard-of-care cisplatin enhanced dendrimer association with 4T1 TNBC cells in vitro

Flow cytometry (Fig. 1A) demonstrated a low but measurable shift in fluorescence intensity of 4T1 TNBC cells upon incubation with Cy5-labeled dendrimer-conjugates, suggesting significant binding. The time-dependent extents of association of Cy5-labeled dendrimer-conjugates with 4T1 cells (Fig. 1B, white symbols), conducted at both 4°C (to assess binding with minimal internalization) and 37°C (to evaluate binding and potential cell internalization) were comparable, suggesting that binding of dendrimer-conjugates to the cell surface was accompanied by limited internalization. Given that cisplatin is a standard-of-care treatment for TNBC, the cell association of Cy5-labeled dendrimer-conjugates was also evaluated in the presence of cisplatin. Dendrimer uptake at 37°C significantly increased when cells were either co-exposed (grey symbols) or pre-exposed (black symbols) to cisplatin, with pre-exposure resulting generally in higher cell association of dendrimers.

**Fig. 1.**
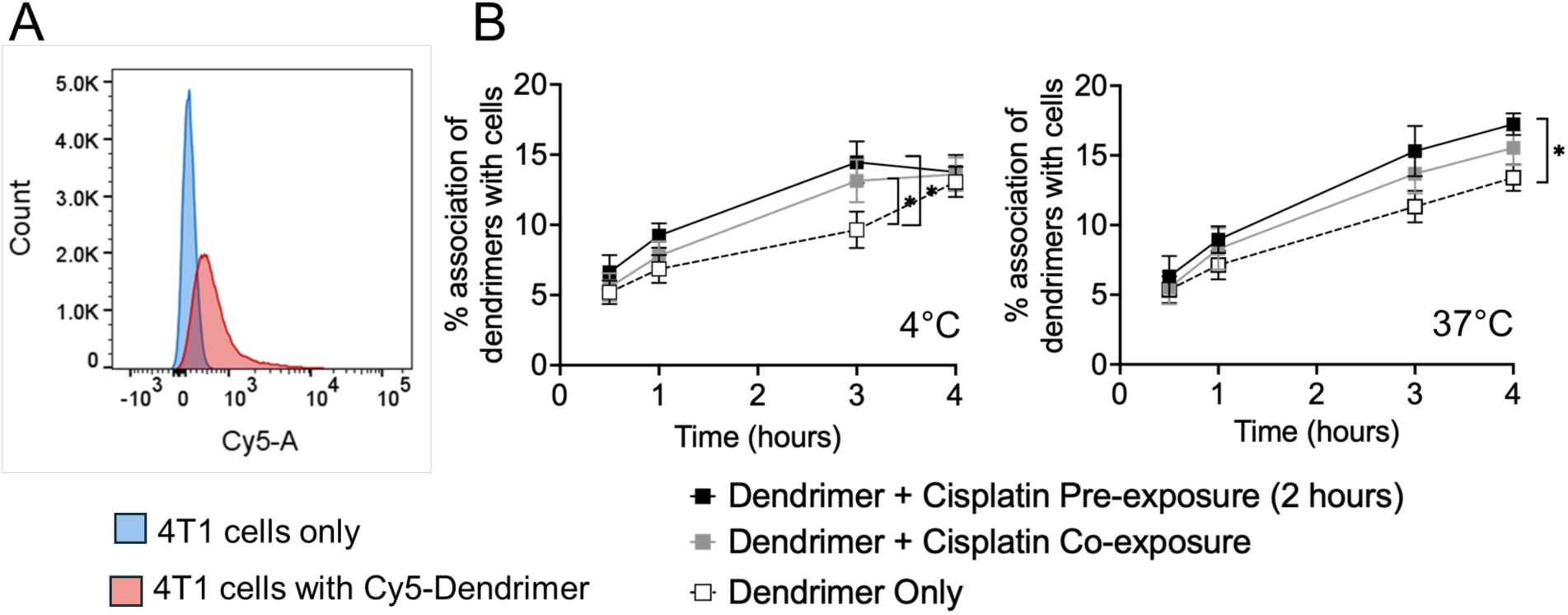
Dendrimers associated with 4T1 murine triple negative breast cancer cells but exhibited limited internalization by cancer cells. (A) Flow cytometry histogram illustrating the association of Cy5-labeled dendrimer-conjugates to 4T1 cells *in vitro*. Cells in suspension (10×10⁶ cells/mL) were incubated with Cy5-labeled dendrimer-conjugates at a dendrimer-to-cell ratio of 50×10⁶:1 for 1 hour on ice. Cells were then washed 3X with cold PBS and were resuspended at a concentration of 1×10⁶ cells/mL before being analyzed by flow cytometry. (B) Extents of time-dependent association of Cy5-labeled dendrimers with 4T1 cells at 4°C (left), that limited endocytosis, and 37°C (right), allowing endocytosis, were comparable. Cells were either pretreated with cisplatin before incubation with Cy5-labeled dendrimer-conjugates (cisplatin; black), concurrently exposed to cisplatin and dendrimer-conjugates (grey symbols), and/or exposed to dendrimer-conjugates alone without cisplatin (white symbols). For uptake experiments, cells in suspension (2×10⁶ cells/mL) were incubated with dendrimer-conjugates (20.4 µg/mL) at a dendrimer-to-cell ratio of 50×10⁶:1. Cisplatin concentration was maintained at (0.4 µg/mL). Data shown are the means ± standard deviation from n = 3 independent experiments. * indicates p-value < 0.05.

### Dendrimer-radioconjugates: enhancement of cell kill in monolayers by low-dose standard-of-care cisplatin

The clonogenic survival of 4T1 cells exposed to actinium-225 dendrimer-conjugates ([^225^Ac]Ac-DOTA-dendrimer) decreased in an activity concentration-dependent manner (white/open symbols, Fig. 2A) and was further lowered when cells were co-treated with low concentrations of cisplatin (10% and 20%, grey and black symbols, respectively, of its IC_50_ (Fig. S1)). The reduction in clonogenic survival when 4T1 cells were treated with both modalities, was partly attributed to the greater association of activity ([^225^Ac]Ac-DOTA-dendrimer) with cells, as illustrated in Fig. 2B. In particular, the survival fraction of cells, that were exposed to 74kBq/mL of [^225^Ac]Ac-DOTA-dendrimer (indicated in blue hues in Fig. 2A), correlated with the cell-associated activity, that increased with increasing cisplatin concentrations in the incubation medium (indicated by the darker blue hues). (This trend was observed independent of the levels of incubation activity of [^225^Ac]Ac-DOTA-dendrimer, as shown in Fig. S2 (A), and independent of the form of activity (the [^225^Ac]Ac-DOTA chelate without the dendrimer resulted in similar findings (Fig. S2 (B)).

**Fig. 2.**
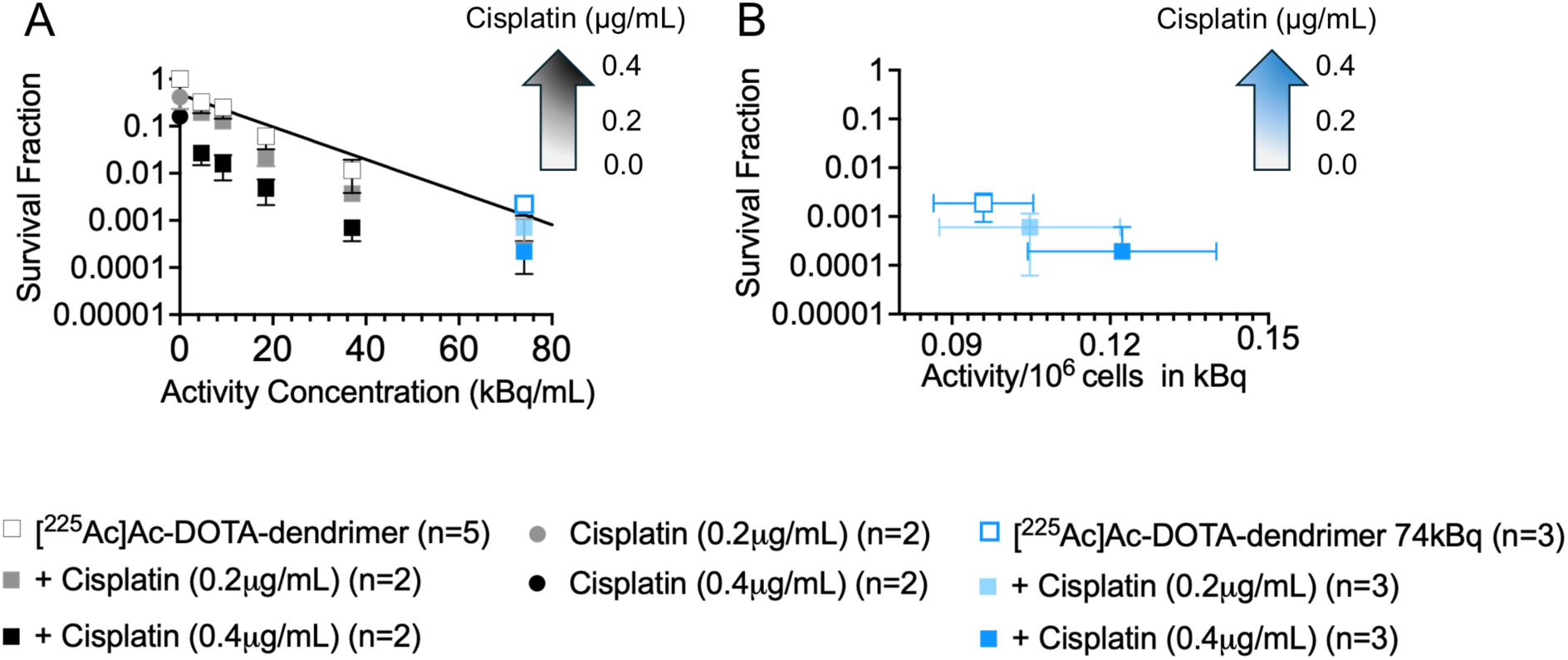
Increasing concentrations of cisplatin enhanced the efficacy of [^225^Ac]Ac-DOTA-dendrimers to inhibit the 4T1 cells’ clonogenic survival (A) and resulted in increased mean activities associated per cell that were correlated with lower clonogenic survival (B). Clonogenic survival fractions of 4T1 cells (A) as a function of the activity concentrations of actinium-225 dendrimer-conjugates ([^225^Ac]Ac-DOTA-dendrimers) alone (white squares) and/or in the presence of cisplatin (0.2/ 0.4 µg/mL; gray / black symbols, respectively); and (B) as a function of cell-associated activity for those cells shown in blue symbols in Fig. 2A that were treated with 74kBq/mL (2μCi/mL). Cells were incubated with treatments for 6 hours. The dendrimer mass concentration was constant at 10µg/mL. Data represent mean ± standard deviation from n = 3 independent experiments.

### Dendrimer-radioconjugates: enhanced spheroid growth inhibition by low-dose standard-of-care cisplatin

In 4T1 multicellular spheroids, employed as surrogates of the tumors’ avascular regions [16], the time-integrated radial distributions of fluorescent dendrimer-conjugates demonstrated their predominant localization at the spheroid periphery (Fig. 3A), with limited penetration in regions deeper than 100 µm (the spatio-temporal distributions in spheroids, that were exposed to dendrimer-radioconjugates for up to 2 hours, are shown in Fig. S3 (A), (B)). This peripheral localization was attributed to the significant binding of dendrimers to 4T1 cells (Fig. 1A) as previously observed with dendrimers and other cancer cells and their spheroids [11]. In contrast to dendrimers’ penetration profile, the time-integrated concentration of CFDA-SE, used as a fluorescent surrogate for cisplatin [15], demonstrated deeper penetration toward the spheroid core region (the spatio-temporal distributions are shown in Fig. S3 (C), (D)). Therefore, cells residing on the spheroid periphery could be potentially exposed to both therapeutic modalities and cells in the spheroid core would be exposed to cisplatin but not be irradiated by α-particles.

**Fig. 3.**
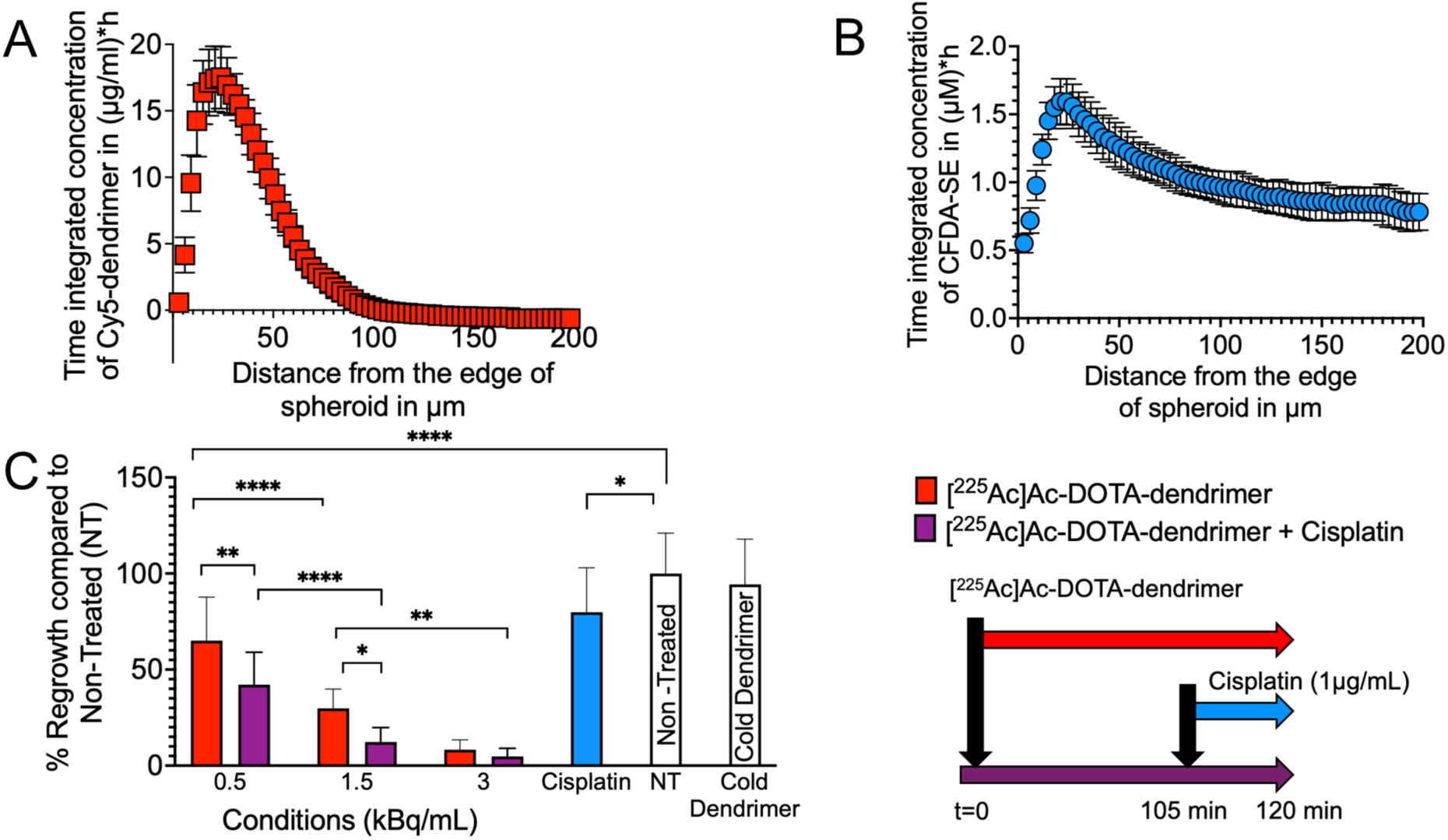
The different microdistributions of fluorescent-dendrimers and of CFDA-SE, a fluorescent surrogate for cisplatin, suggested that cells in spheroids were collectively exposed either to both modalities (cells in the spheroid periphery/edge) or at least to cisplatin (cells in the spheroid core), and, when the two modalities (dendrimer-radioconjugates and cisplatin) acted together on spheroids, they resulted in best inhibition of spheroid regrowth compared to any single modality treatment. Time-integrated radial microdistributions in 4T1 spheroids of (A) Cy5-labeled dendrimer-conjugates, and (B) CFDA-SE. Spatial profiles of each agent vs. time are shown in Fig. S3 (A-D). (C) The extent of inhibiting spheroid regrowth (assay used as surrogate of tumor recurrence) of 400µm-in-diameter 4T1 spheroids treated with [^225^Ac]Ac-DOTA-dendrimer (red bars), cisplatin (blue bar) and/or their combination (purple bars) relative to untreated spheroids and the cold dendrimer. (D) Treatment schedule: Spheroids were exposed to dendrimer-radioconjugates for 2 hours in the absence or presence of cisplatin (0.25 hours), and/or to cisplatin alone (0.25 hours). Incubation times were chosen to approximately scale with the relevant blood circulation kinetics of each modality. Mean values ± the standard deviations of n=6 spheroids per condition (n=3 independent dendrimer-radioconjugate preparations) are shown. *** indicates p-value <0.001; **<0.01; *<0.05.

The effect of combined modalities on reducing the extent of spheroid regrowth was compared to the effect of the dendrimer-radioconjugate alone for different activity concentrations. At 0.5kBq/mL the mean difference in spheroid regrowth with and without cisplatin was 22.86% (p-value = 0.0031), at 1.5kBq/mL was 17.95% (p-value = 0.0459), and at 3.0kBq/mL was not significant (3.50%, p-value = 0.9987). The cold dendrimer did not affect the extent of spheroid regrowth compared to untreated groups, confirming that dendrimers alone were non-toxic at the concentrations employed in this study. The microdistribution patterns shown in Fig. 3A and 3B partly explained the findings from the spheroid regrowth assays (Fig. 3C), where the combined treatment of actinium-225 dendrimer-conjugates and cisplatin exhibited *superior inhibition* of spheroid regrowth compared to *either treatment modality alone*. At the spheroid periphery, where usually more aggressively growing cells resided, localization of both modalities potentially formed a more lethal cocktail (in agreement with Fig. 2A). Additionally, in the deeper spheroid regions, which were not expected to be reached and/or be irradiated by the dendrimer-radioconjugates, cisplatin penetration (as implied by its surrogate’s spatiotemporal profiles, Fig. 3B) potentially decreased cell survival.

### Dosimetry and long-term toxicities in mice

The pharmacokinetics of [¹¹¹In]In-DTPA-dendrimer, with and without pretreatment with cisplatin, on the three tumor-bearing mouse models are shown in the SI section (Fig. S4, and Tables S1-S6) and were employed to calculate the dosimetry of [^225^Ac]Ac-DOTA-dendrimer. In all mouse models, the tumor absorbed doses of actinium-225 delivered by dendrimers were selectively increased with low-dose cisplatin pretreatment, that was administered one day before the systemic administration of dendrimer-radioconjugates (Table 2 and Fig. 4). The extents of tumor absorbed dose increase depended on the tumor’s implantation site and were: 1.5 x greater (p-value = 0.0023), 1.6 x greater (p-value = 0.0024), and/or 1.8 x greater (p-value = 0.1275), for the subcutaneous, orthotopic/mammary fat pad, and intracranial sites, respectively. Interestingly, the normal-organ dosimetry was unaffected by cisplatin pretreatment, in agreement with previous reports on an orthotopic glioblastoma mouse model (*8*).

**Fig. 4.**
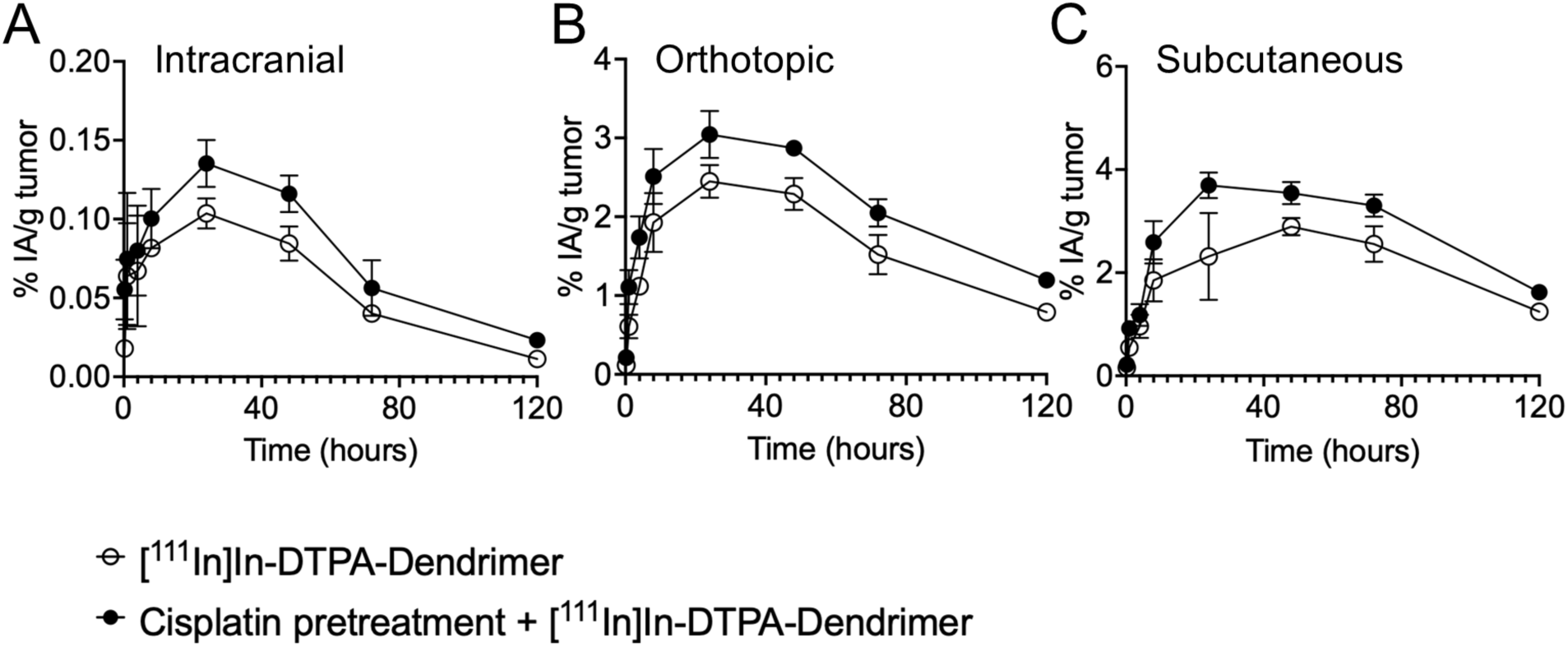
Pretreatment with cisplatin selectively enhanced the tumor uptake of systemically injected [¹¹¹In]In-DTPA-dendrimer in Balb/c mice, irrespective of the implanted 4T1 tumor anatomic sites. The normal-organ uptake was unaffected by the administration of cisplatin (see Table 2, Fig. S4). At specified time points, mice were euthanized, and tumors along with selected tissues were harvested, weighed, and measured for radioactivity using gamma counting. Data are presented as decay-corrected percent injected activity per gram of tumor tissue (% IA/g).

**Table 2.** Tumor absorbed doses delivered by systemically injected [²²⁵Ac]Ac-DOTA-dendrimer were selectively increased with low-dose cisplatin pretreatment, injected one day earlier, independent of tumor implantation/anatomic site. Normal-organ dosimetry was unaffected by cisplatin pretreatment. Dosimetry calculations for [²²⁵Ac]Ac-DOTA-dendrimer using the biodistributions of systemically administered [¹¹¹In]In-DTPA-dendrimer alone or with cisplatin pre-treatment (24 hours earlier) (see Tables S1-S6). Cisplatin was administered intraperitoneally (5mg/kg mouse) to enable its relatively slow clearance in the blood and to imitate its long(er) infusions in the clinic. Dosimetry calculations were based on the same administered activities employed in the corresponding therapeutic assessment studies: namely 25.9kBq and 22.2kBq of [²²⁵Ac]Ac-DOTA-dendrimer per 18 g mouse for the intracranial tumor models and the orthotopic/mammary fat-pad (and subcutaneous) tumor models, respectively, using the software package RAPID Dosimetry (Baltimore, MD). All daughters’ (α-particle and electron) energy delivered to tumors by dendrimers was assumed to be absorbed by the tumor, as previously reported (*8*). Mean absorbed doses ± standard deviations are reported (n=2 mice per time point). * and ǂ indicate *p*-values<0.01.

| Organ | Absorbed Dose (Gy) |  |  |  |  |  |
| --- | --- | --- | --- | --- | --- | --- |
|  | Intracranial |  | Orthotopic |  | Subcutaneous |  |
|  | Without cisplatin | With cisplatin pretreatment | Without cisplatin | With cisplatin pretreatment | Without cisplatin | With cisplatin pretreatment |
| Tumor | 0.17 ± 0.02 | 0.30 ± 0.07 | 2.36 ± 0.06* | 3.81 ± 0.08* | 4.31 ± 0.06† | 6.54 ± 0.14† |
| Brain | 0.05 ± 0.00 | 0.07 ± 0.00 | 0.03 ± 0.00 | 0.05 ± 0.00 | 0.06 ± 0.01 | 0.02 ± 0.00 |
| Heart | 0.77 ± 0.05 | 0.76 ± 0.18 | 0.42 ± 0.02 | 0.99 ± 0.14 | 0.65 ± 0.05 | 0.57 ± 0.02 |
| Lungs | 0.00 ± 0.00 | 0.00 ± 0.00 | 0.00 ± 0.00 | 0.00 ± 0.00 | 0.00 ± 0.00 | 0.00 ± 0.00 |
| Stomach | 0.00 ± 0.00 | 0.00 ± 0.00 | 0.00 ± 0.00 | 0.00 ± 0.00 | 0.00 ± 0.00 | 0.00 ± 0.00 |
| Liver | 0.24 ± 0.01 | 0.36 ± 0.01 | 0.09 ± 0.00 | 0.11 ± 0.01 | 0.21 ± 0.01 | 0.34 ± 0.02 |
| Spleen | 0.08 ± 0.01 | 0.16 ± 0.00 | 0.04 ± 0.00 | 0.06 ± 0.00 | 0.05 ± 0.00 | 0.05 ± 0.00 |
| Intestines | 0.00 ± 0.00 | 0.00 ± 0.00 | 0.00 ± 0.00 | 0.00 ± 0.00 | 0.00 ± 0.00 | 0.00 ± 0.00 |
| Kidneys | 0.94 ± 0.05 | 1.24 ± 0.09 | 0.71 ± 0.02 | 0.76 ± 0.02 | 0.90 ± 0.02 | 1.17 ± 0.03 |

In Balb/c mice implanted with 4T1 tumors intracranially, orthotopically and/or subcutaneously, the tumor absorbed doses, with or without pretreatment with cisplatin, followed the order: subcutaneous tumors > orthotopic tumors > intracranial tumors (Table 2 and Fig. 4).

Long-term toxicity studies of systemically injected [²²⁵Ac]Ac-DOTA-dendrimers were conducted over a 10-month period in tumor-free Balb/c mice at administered activities of 25.9kBq (700nCi), 33.3kBq (900nCi) or 51.8kBq (1400nCi) per 18g mouse, both with and without pretreatment with cisplatin (5mg/Kg). All animals survived during the 10-month period. No long-term pathological concerns were observed in the brain, heart, and lungs at any tested dose in surviving animals (Fig. S5). At 25.0kBq injected activity, no significant toxicities were observed. The group that was treated with 33.3kBq TAT, after 10 months, exhibited mildly enlarged hepatocytes and mildly enlarged renal tubular epithelial cells with karyomegaly in both cell types (Fig. 5). The spleen also demonstrated mild reduction in white pulp, suggesting atrophy or previous cell death. However, at 51.8kBq, notable long-term toxicities were observed and included renal tubular atrophy, dilated tubules, mild single cell death, and renal tubular karyomegaly (indicated by the arrow and the black squared region in the kidneys, Fig. 5 E, F). The liver also displayed karyomegaly with enlarged hepatocytes and nuclei. The spleen showed further white pulp reduction. Based on these findings, 51.8kBq was considered a toxic dose for Balb/c mice (18g) over a 10-month period.

**Fig. 5.**
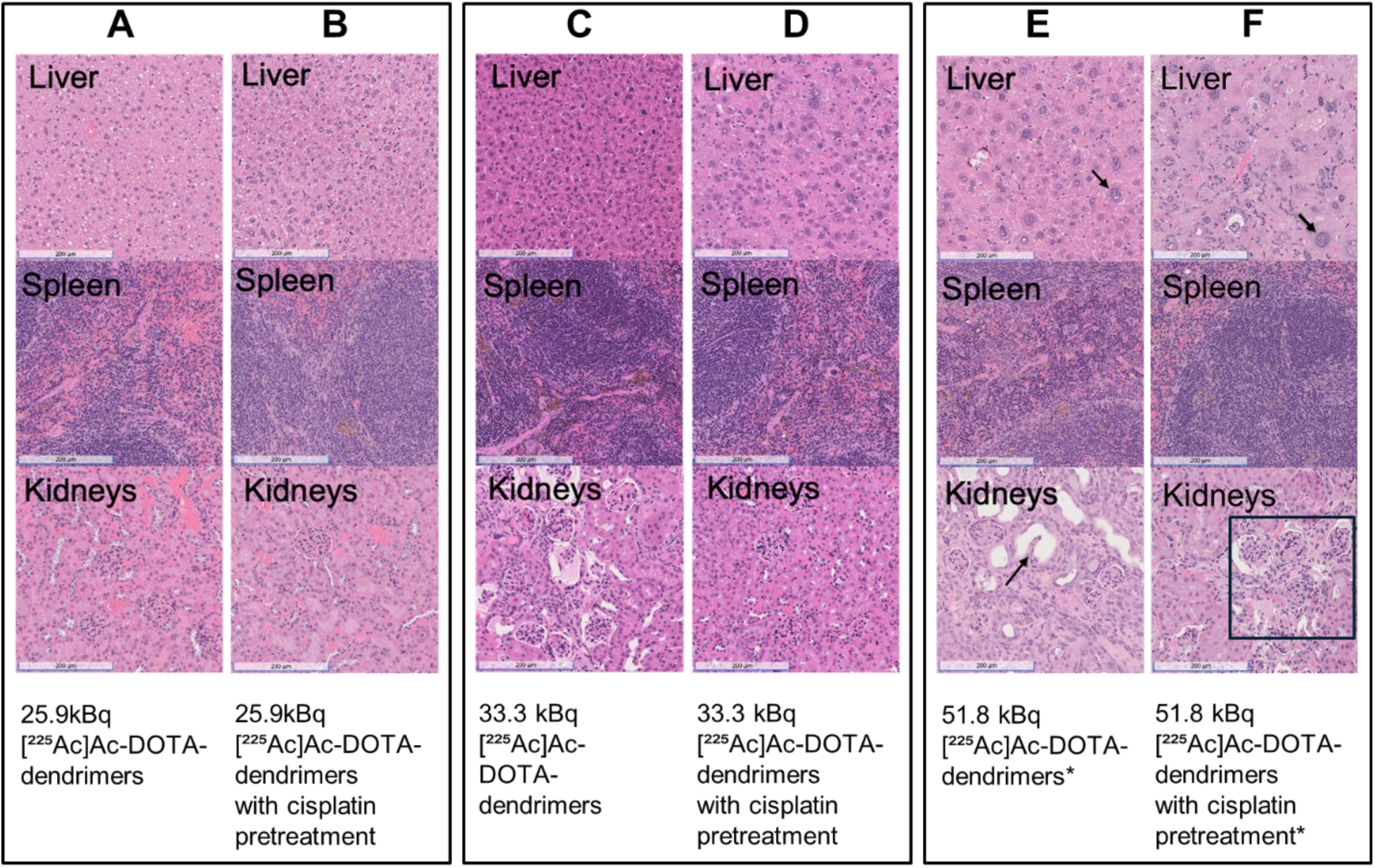
Acute toxicities in Balb/c tumor-free mice were not observed at any of the systemically injected activities of [^225^Ac]Ac-dendrimers (with and without pretreatment with low-dose cisplatin). The onset of long-term toxicities (10-months after injection) was observed by histopathology only at the highest systemically injected activities of 51.8 kBq (indicated by the arrow and the black squared region in the kidneys). The mice didn’t show any physical signs of toxicity at any level of injected activities. Six-week-old mice were treated with 25.9, 33.3 and/or 51.8kBq [²²⁵Ac]Ac-DOTA-dendrimers/18g mouse, with or without cisplatin pretreatment (5mg/Kg) administered 24 hours earlier. During the 10-month observation period, none of the treated mice reached any of the study’s endpoints. Ten months after administration of activity, mice were euthanized, and organs were harvested for histopathological analysis using H&E staining. The scale bar corresponds to 200μm on all images (H&E of the brain, heart and lungs are shown in Fig. S5).

### Therapeutic efficacy of actinium-225 dendrimer-radioconjugates against 4T1 tumors implanted at different anatomic sites

In treatment studies, mice were systemically injected with [^225^Ac]Ac-DOTA-dendrimers at 25.9kBq per 18g mouse (in the intracranial tumor model) or 22.2kBq per 18g mouse (in the orthotopic and subcutaneous models), either as monotherapy or 24 hours after cisplatin pretreatment (5mg/Kg). Irrespective of the tumor implantation site, actinium-225 delivered by dendrimer-radioconjugates significantly prolonged survival in all mouse models when injected a day after cisplatin pretreatment (Fig. 6A, 7A, 8A). In particular, in the intracranial tumor model (Fig. 6A), the mean survival of mice treated with both agents (14 days) was significantly longer than the mean survival of mice treated with TAT alone (9 days, *p*-value = 0.0039), cisplatin alone (12 days, *p*-value = 0.0321), and/or not being treated (10 days, *p*-value = 0.0476). In the orthotopic tumor model (Fig. 7A), the mean survival of mice treated with both agents (18 days) was significantly longer than the mean survival of mice treated with TAT alone (13 days, *p*-value = 0.0039), chemotherapy alone (13 days, *p*-value = 0.0072), and/or not being treated (13 days, *p*-value = 0.0006). And, similarly, the subcutaneous tumor model (Fig. 8A), exhibited the largest difference among cohorts (that correlated with the greatest tumor absorbed doses among all three tumor sites (Table 2)), with the mean survival of mice treated with both agents (24 days) being significantly longer than the mean survival of mice treated with TAT alone (17 days, *p*-value <0.0001), cisplatin alone (16 days, *p*-value = 0.0001), and/or not being treated (15 days, *p*-value = <0.0001). With the exception of the mouse model with intracranial 4T1 tumors, which did not generate adequate contrast to be imaged by MRI, longer survival correlated with slower tumor growth (Fig. 7B, 8B).

**Fig. 6.**
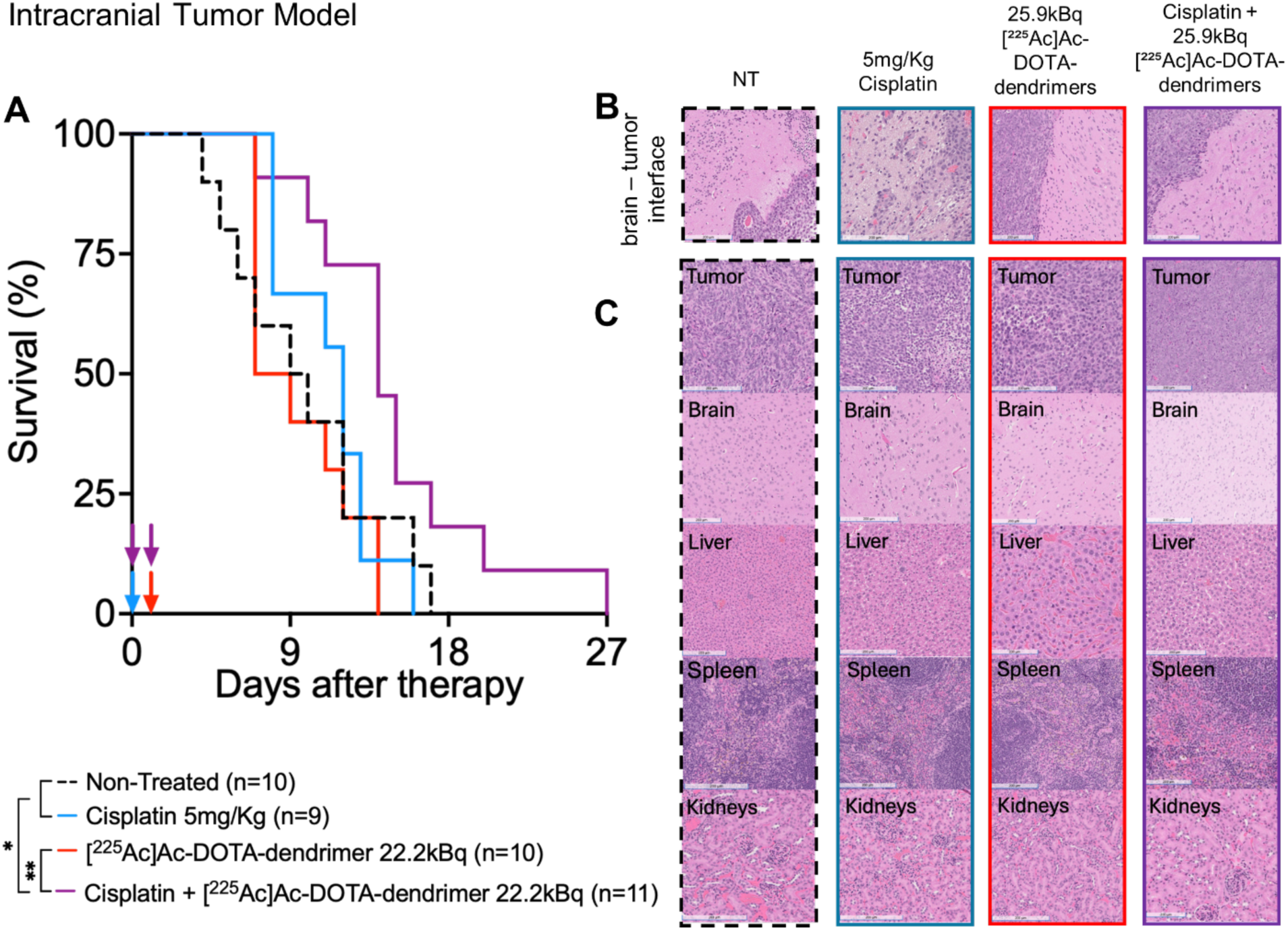
Balb/c mice with syngeneic 4T1 triple negative breast cancer implanted intracranially: Combining [^225^Ac]Ac-DOTA-dendrimers and low-dose cisplatin pretreatment significantly prolonged survival compared to either modality alone, and did not affect the nearby healthy brain. (A) Survival of 4T1 tumor-bearing Balb/c mice treated with intravenous injection of [^225^Ac]Ac-DOTA-dendrimers (25.9kBq per 18g mouse on day 1 as shown, or day 10 after tumor inoculation), with intraperitoneal injection of 5mg cisplatin/Kg mouse (on day 0), and/or their combination (cisplatin on day 0, αRPT on day 1) relative to non-treated mice. Vertical arrows indicate treatment scheduling. The study endpoint was determined by weight loss ≥20% (shown in Fig. S6). Statistical significance was evaluated by the log-rank test. ** indicates *p*-value <0.01; (B) Characteristic H&E-stained sections of the tumor-healthy brain interface indicating lack of adverse effects on the tumor-neighboring brain (scale bar = 200µm). (C) H&E-stained sections of tumors and normal organs of mice sacrificed after indicated treatment (see also Fig. S6 for H&E of lungs and heart). No significant renal, hepatic or splenic toxicity was observed. Scale bar=200µm.

**Fig. 7.**
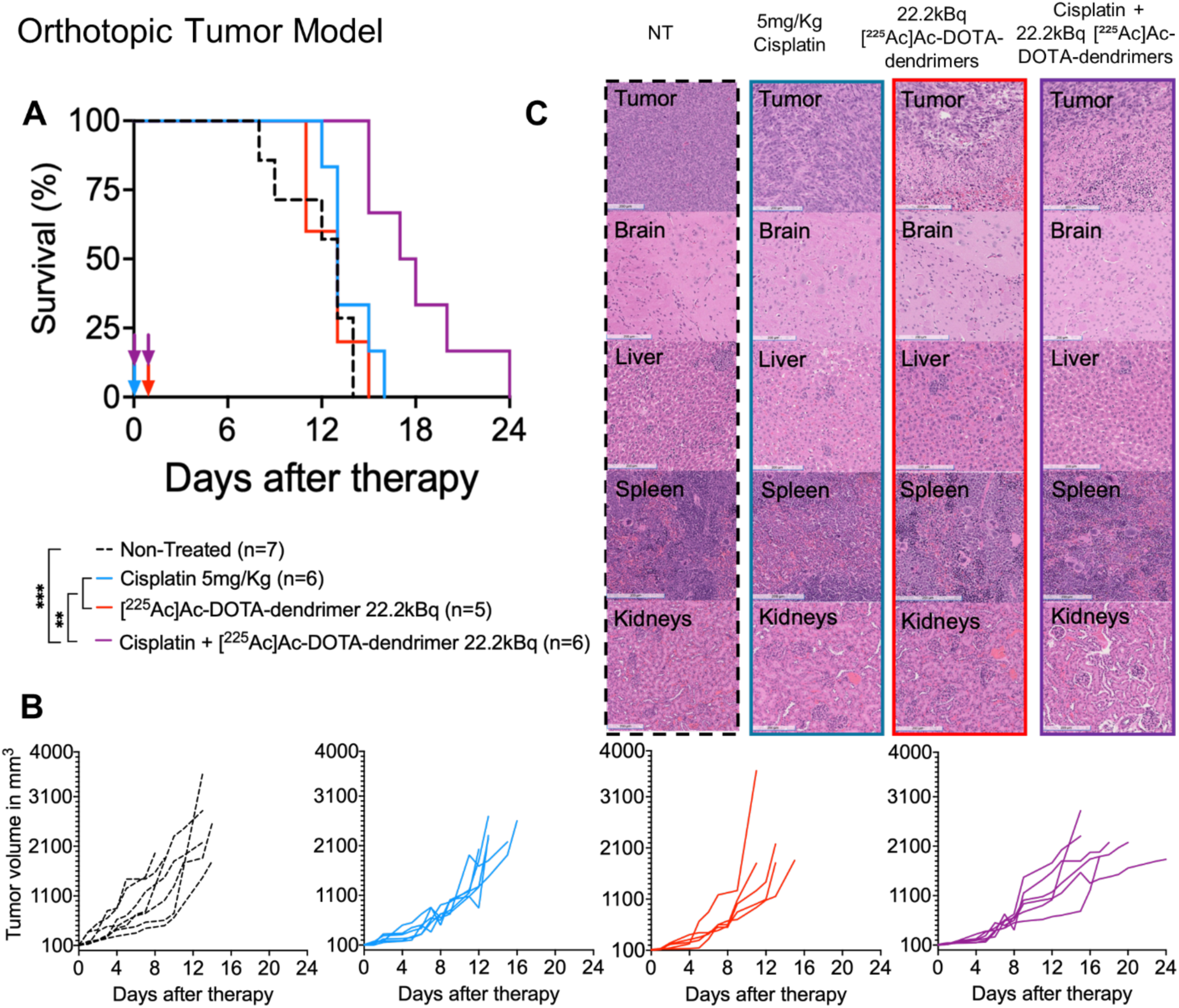
Balb/c mice with syngeneic 4T1 triple negative breast cancer implanted in the mammary fat pad: Combining [^225^Ac]Ac-DOTA-dendrimers and low-dose cisplatin pretreatment significantly prolonged survival compared to either modality alone due to better tumor growth inhibition. (A) Survival of 4T1 tumor-bearing Balb/c mice treated with intravenous injection of [^225^Ac]Ac-DOTA-dendrimers (22.2kBq per 18g mouse on day 1), with intraperitoneal injection of 5mg cisplatin/Kg mouse (on day 0), and/or their combination (cisplatin on day 0, αRPT on day 1) relative to non-treated mice. Treatment was initiated when tumors reached 100mm^3^. Vertical arrows indicate treatment scheduling. The study endpoint was determined either by weight loss ≥20% (shown in Fig. S7), or by the tumor weight becoming greater than 10% of body weight at the day of injection of therapy, or by the tumor interfering with the movement or function of mice. Statistical significance was evaluated by the log-rank test. ** indicates *p*-value <0.01; *** <0.001. (B) Individual tumor volumes over time for each treatment group. (C) Characteristic H&E-stained sections of tumors and normal organs of mice sacrificed after indicated treatment (see also Fig. S7 for H&E of lungs and heart). No significant renal, hepatic or splenic toxicity was observed. Scale bar = 200µm.

**Fig. 8.**
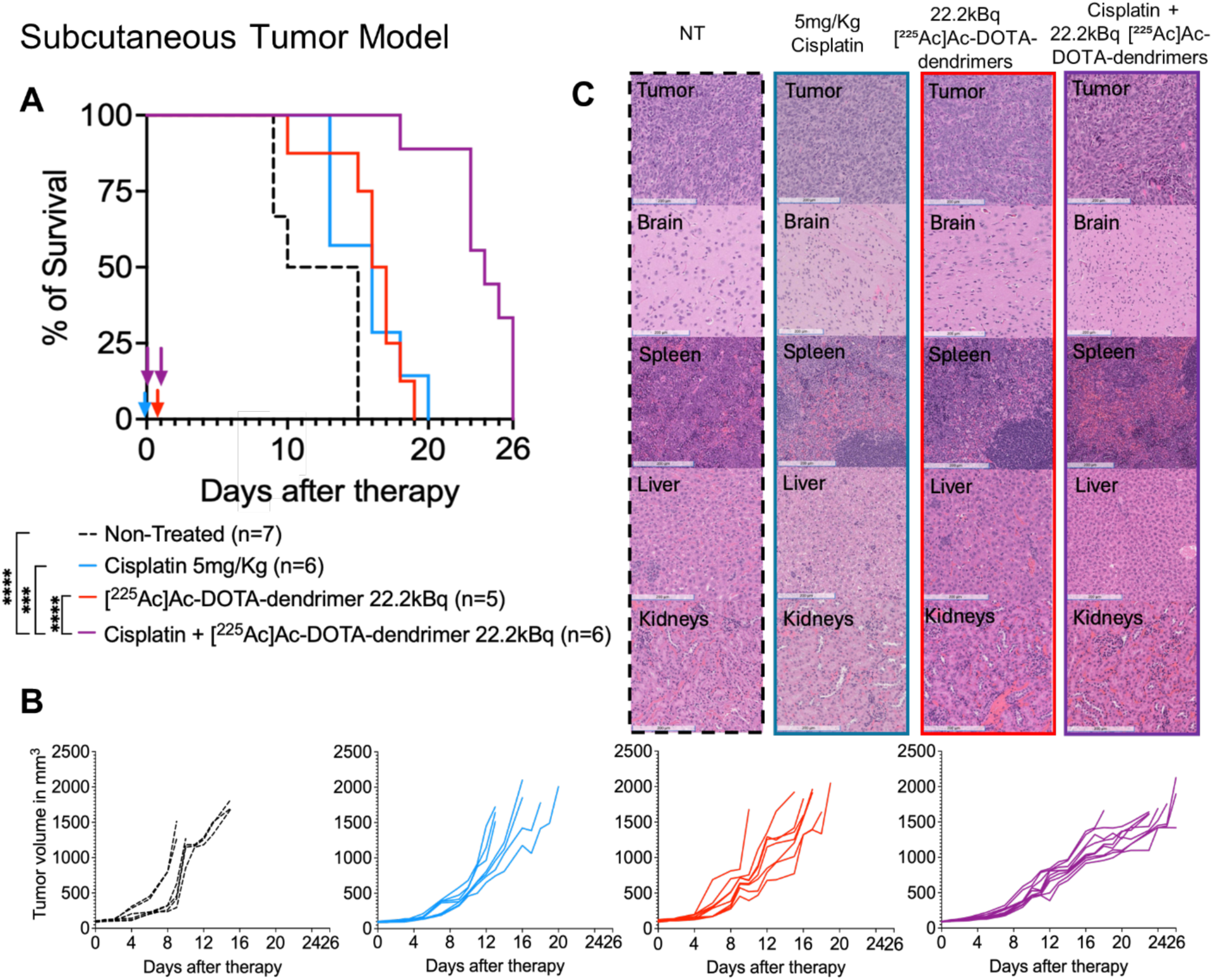
Balb/c mice with syngeneic 4T1 triple negative breast cancer implanted subcutaneously: Combining [^225^Ac]Ac-DOTA-dendrimers and low-dose cisplatin pretreatment significantly prolonged the survival compared to either modality alone due to better tumor growth inhibition. (A) Survival of 4T1 tumor-bearing Balb/c mice treated with intravenous injection of [^225^Ac]Ac-DOTA-dendrimers (22.2kBq per 18g mouse on day 1), with intraperitoneal injection of 5mg cisplatin/Kg mouse (on day 0), and/or their combination (cisplatin on day 0, αRPT on day 1) relative to non-treated mice. Treatment was initiated when tumors reached 100mm^3^. Vertical arrows indicate treatment scheduling. The study endpoint was determined either by weight loss ≥20% (shown in Fig. S8), or by the tumor weight becoming greater than 10% of body weight at the day of injection of therapy, or by the tumor interfering with the movement or function of mice. Statistical significance was evaluated by the log-rank test. ** indicates *p*-value <0.01; **** <0.0001. (B) Individual tumor volumes over time for each treatment group. (C) Characteristic H&E-stained sections of tumors and normal organs of mice sacrificed after indicated treatment (see also Fig. S8 for H&E of lungs and heart). No significant renal, hepatic or splenic toxicity was observed. Scale bar = 200µm.

At the injected activities used, that did not cause any long-term toxicities (Fig. 5), TAT as monotherapy did not result in significant survival advantage relative to non-treated mice with intracranial or orthotopic tumors, possibly due to the corresponding relatively low tumor absorbed doses (Table 2).

Importantly, in the intracranial tumor model, the pathologic evaluation of the tumor-brain interface across all treated groups revealed no adverse effects by TAT or by cisplatin on the surrounding healthy brain tissue (Fig. 6B). Regarding toxicities in general, in all tumor-bearing mice, at the time of sacrifice (when either of the end points were reached), pathology assessment of excised tissues did not indicate noteworthy renal, hepatic, or splenic toxicities in any treated group (Fig. 6C, 7C, 8C and Fig. S6, S7, S8). Mild to moderate levels of extramedullary hematopoiesis (EMH) were observed in the liver and spleen across all three tumor models (intracranial, orthotopic, and subcutaneous). In the subcutaneous model, the non-treated group exhibited the highest EMH levels in the liver, while EMH was reduced in treated conditions, with the lowest levels observed in the groups that received the combined modalities (TAT with cisplatin pretreatment) and correlated with the tumor burden at the time of sacrifice (all mouse weight measurements over time are plotted in Fig. S9-S11).

## Discussion

To selectively and effectively kill TNBC cells by targeted alpha-particle therapy (TAT), a hydroxyl-terminated, generation 6, DOTA-functionalized PAMAM dendrimer nanoparticle was employed to deliver the α-particle emitter actinium-225. The established systemic targeting of tumors, even in the brain, enabled by this clinically-validated platform was utilized to improve efficacy and reduce side effects of TAT, synergistically with cisplatin. *In vitro*, on cell monolayers and in spheroids, the ability of dendrimer-radioconjugates to extensively associate and effectively kill murine 4T1 TNBC cells was confirmed and was shown to be enhanced by low concentrations of standard-of-care cisplatin. *In vivo*, systemically injected actinium-225 dendrimer-radioconjugates: (1) demonstrated selective tumor uptake, independent of the implanted tumors’ anatomic site; (2) exhibited slow clearance from tumors, increasing the tumor absorbed dose; and (3) significantly prolonged survival of mice, when combined with low-dose cisplatin pretreatment, in a proof-of-concept study on mice with syngeneic 4T1 TNBC tumors implanted orthotopically or intracranially or subcutaneously, as model sites of recurrence and/or metastasis.

Prolonged survival of mice treated with the combined modalities was correlated with greater tumor growth inhibition (on the orthotopic and subcutaneous models, the two cases we could measure the tumor growth). This finding was attributed to three factors. First, to the greater cell killing efficacy when both modalities acted on the same cancer cells, as observed *in vitro* on cell monolayers. Second, investigation of the spatiotemporal distributions of dendrimer-conjugates and cisplatin surrogates on 4T1 spheroids, employed to capture the diffusion-limited environment of the tumors’ avascular regions, suggested two points: (a) that the improved killing efficacy of the combined modalities was not only due to both agents acting on the same population of cells (cells that resided closer to the spheroids’ periphery, which represented areas corresponding to the perivascular regions in solid tumors), but (b) that the improved efficacy was additionally attributed to the killing of cells residing in the spheroid core (which represented areas far from the tumor vasculature in solid tumors), which were shown to be exposed to cisplatin and that would otherwise probably not be significantly affected by the dendrimer-delivered TAT, due to its limited penetration. Third, in agreement with a previous report on an orthotopic glioblastoma mouse model, pretreatment of mice with low dose chemotherapy resulted in selective increase in tumor uptake of dendrimer-radioconjugates without affecting the normal organ uptake. The increased tumor absorbed doses with cisplatin pretreatment seem to be a yet another reason for the combined modalities conferring significant survival advantage relative to each modality alone.

Dosimetry evaluation on all three mouse models suggested that renal toxicities could act as the dose limiting factor given that dendrimers were shown to clear through the kidneys, both in mice and in humans [10, 12, 17, 18]. Notably, however, this study demonstrated significant therapeutic advantage at dendrimer-delivered TAT activities that were shown to not cause any long-term (10-months) toxicities, including at the kidneys.

To potentially decipher any potential role of the tumor’s anatomic site on the treatment response of the different mouse models, the mean survival (from the day of treatment initiation) was plotted as a function of the tumor absorbed dose delivered by actinium-225 dendrimer-radioconjugates with and without cisplatin pretreatment. Fig. 9 demonstrates a strong unified correlation, largely independent of the tumor anatomic site, with the exception of intracranial tumors exposed to cisplatin pretreatment. Although the studies herein were performed in syngeneic mouse models with implanted, rather than spontaneous, tumors at these different sites, in terms of delivery potential, these findings could be supportive of the dendrimer-radioconjugate as a single type of TAT, that could be a useful therapeutic tool to simultaneously treat multi-site TNBC metastases, including TNBC metastases in the brain. A major delivery obstacle to tumors in the brain is the limited tumor vascular permeability, and these dendrimers seem to be ideal to permeate the blood brain tumor barrier and deliver to intracranial tumors lethal doses that are further selectively enhanced by low dose standard-of-care cisplatin.

**Fig. 9.**
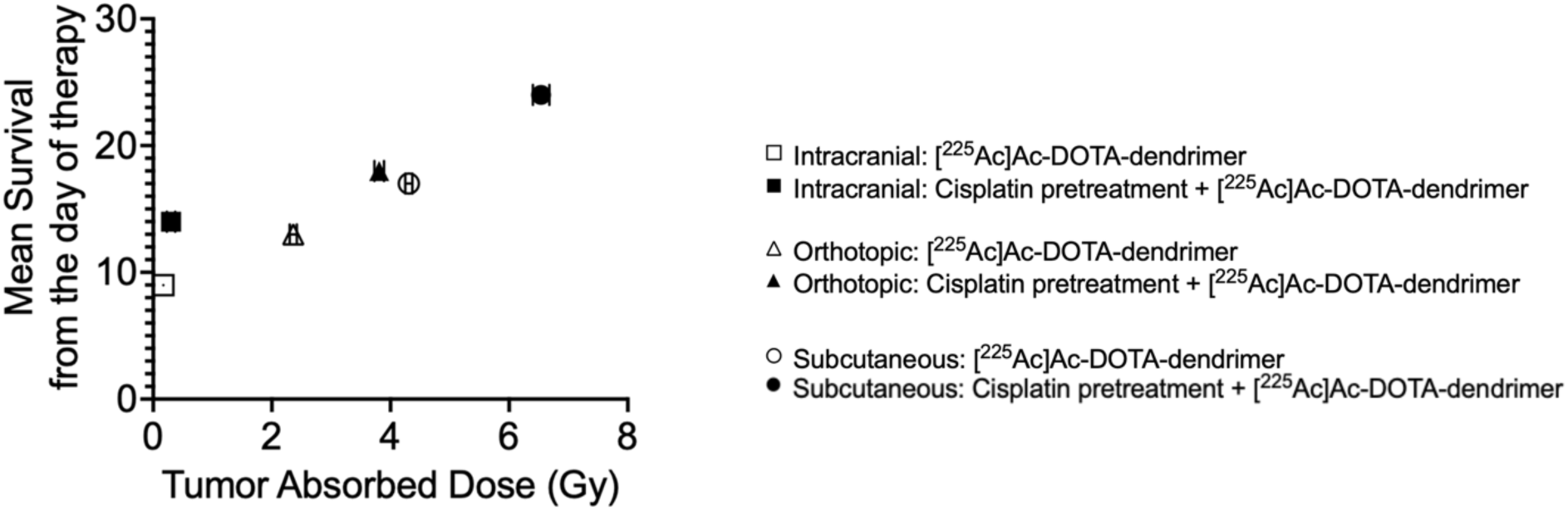
Mean survival of tumor-bearing mice treated with the combined modalities (shown in Fig 6A, 7A and 8A) correlated strongly with the tumor absorbed doses (Table 2) and appeared independent of the implanted tumors’ anatomic site.

The response of mice bearing tumors simultaneously at all three anatomic sites were not evaluated, but it is highly possible that the levels of tumor growth inhibition in such a multi-site model would be similar to the studies herein, where a single tumor anatomic site was included per mouse model. The main reason for this should be the low fractions of tumor delivered doses in all cases studied, which would not be expected to affect/limit the absorbed doses to additional tumors in the same mouse. However, the effect of a multi-site tumor burden on the animals’ survival could be a factor introducing additional complications and less tolerance to injected activities. We kept the injected TAT activities, low, yet but effective, by including pretreatment with low-dose standard-of-care cisplatin, to potentially preserve the safety of this approach.

## Conclusion

Patients with advanced TNBC develop metastatic soft-tissue solid tumors usually in the brain, the lungs and/or liver. A single carrier, that delivers lethal TAT doses to all these different anatomic sites, could be useful in the clinic. In this study, the potential of a systemically injected dendrimer-delivered TAT, combined with low-dose standard-of-care chemotherapy, to safely inhibit tumor growth, was demonstrated against TNBC tumors in the brain, subcutaneously (as a general, well-vascularized soft-tissue tumor surrogate), and orthotopically (as a surrogate of a recurrent tumor).

## Acknowledgements

The authors thank Dr. Ines Godet, Natalie Joe, and Dr. Daniele Gilkes for providing the murine cell line, Dr Aprameya Prasad for support with the establishment of the animal models, Dr. George Sgouros for support with the dosimetry calculations, and Dr. Laura Ensign for support with the intracranial mouse studies.

## Author Contributions

Radioconjugate preparation, data collection and data analysis were performed by Pooja Hariharan, Rajiv Nair, and Aira Sarkar. Dendrimer synthesis was performed by Tony Wu, Chang Liu, and Wathsala Liyanage with the guidance of Rangaramanujam Kannan. Pathology evaluation was by Kathleen Gabrielson. The first draft of the manuscript was written by Stavroula Sofou and all authors commented on the manuscript. All authors read and approved the final manuscript.

## Data Availability

The datasets generated during and/or analyzed during the current study are available from the corresponding author on reasonable request.

## Ethics approval

All applicable international, national, and/or institutional guidelines for the care and use of animals were followed.

