## Supplementary material for "Actinium-225 dendrimer-radioconjugates combined with low-dose standard-of-care chemotherapy: site-independent treatment of triple negative breast cancer metastases": Supplementary Material.pdf

**Supporting Information**

**Materials**

Ethylenediaminetetraacetic acid (EDTA) was purchased from Fisher Scientific (Pittsburgh, PA, USA), and Phosphate Buffered Saline (PBS) was purchased from Sigma-Aldrich (Atlanta, GA, USA). The 3kDa MWCO Amicon Ultra-0.5 Centrifugal Filter Unit (Cat. No. UFC5003) was purchased from Milli Pore Sigma (Saint Louis, MO, USA). Matrigel™, Trypsin and the ultra-low adhesion U-shaped 96-well plates (Cat. No.: 7007) were purchased from Corning (Corning, NY, USA), Roswell Park Memorial Institute (RPMI) medium were from ATCC (Manassas, VA, USA), the Fetal Bovine Serum (FBS) was from Omega Scientific (Tarzana, CA, USA) and penicillin-streptomycin were from ThermoFisher Scientific (Waltham, MA, USA). Cyanine 5 (Cy5) was from GE Healthcare Life Science (Pittsburgh, PA). Cisplatin was purchased from Sigma-Aldrich Chemicals (Atlanta, GA). The 3-(4,5-dimethylthiazol-2-yl)-2,5-diphenyltetrazolium bromide (MTT) assay kit was purchased from Promega (Madison, WI, USA), Chelex® resin from Bio-Rad (Hercules, CA, USA), syringe filters (0.22µm, Cat No. 76479-024) from VWR (Radnor, PA, USA).

S-2-(4-Isothiocyanatobenzyl)-diethylenetriamine pentaacetic acid (DTPA-SCN) and S-2-(4-isothiocyanatobenzyl)-1,4,7,10-tetraazacyclododecane-1,4,7,10-tetraacetic acid (DOTA-SCN) were from Macrocyclics (Dallas, TX, USA). Indium-111 ($^{111}$In), indium chloride, was purchased from BWXT (Ontario, Canada). Actinium-225 ($^{225}$Ac), actinium chloride, was supplied by the U.S. Department of Energy Isotope Program, managed by the Office of Science for Nuclear Physics. Trypsin and Matrigel™ were purchased from Corning (Corning, NY, USA). 6 well plates, 48 well plate, and tissue culture dishes were purchased from VWR (Philadelphia, PA, USA) CFDA-SE was purchased from ThermoFisher Scientific (Waltham, MA, USA).

**Table S1**. Decay-corrected % injected activities (%IA) per mass of tissue for 100µL of 740kBq [$^{111}$In]In-DTPA-dendrimers on BALB/c mice with **intracranial 4T1** Triple Negative Breast Cancer. Error bars correspond to standard deviations of n=2 mice per condition per time point.

| % IA [$^{111}$In]In-DTPA-dendrimer | | | | | | | | |
|---|---|---|---|---|---|---|---|---|
| Time (hours) | 0.25 | 1 | 4 | 8 | 24 | 48 | 72 | 120 |
| Tumor | 0.02 ± 0.00 | 0.06 ± 0.03 | 0.07 ± 0.04 | 0.08 ± 0.00 | 0.10 ± 0.01 | 0.08 ± 0.01 | 0.04 ± 0.00 | 0.01 ± 0.00 |
| Brain | 0.02 ± 0.01 | 0.03 ± 0.00 | 0.03 ± 0.01 | 0.06 ± 0.01 | 0.04 ± 0.01 | 0.03 ± 0.01 | 0.02 ± 0.00 | 0.01 ± 0.00 |
| Blood | 49.78 ± 2.78 | 29.46 ± 3.04 | 21.46 ± 2.34 | 13.43 ± 1.41 | 3.45 ± 1.07 | 1.94 ± 0.89 | 1.47 ± 0.45 | 0.12 ± 0.01 |
| Heart | 1.50 ± 0.14 | 0.78 ± 0.08 | 0.50 ± 0.11 | 0.31 ± 0.06 | 0.25 ± 0.07 | 0.09 ± 0.04 | 0.09 ± 0.02 | 0.06 ± 0.03 |
| Lungs | 0.02 ± 0.00 | 0.02 ± 0.00 | 0.02 ± 0.00 | 0.02 ± 0.00 | 0.02 ± 0.00 | 0.02 ± 0.01 | 0.01 ± 0.00 | 0.01 ± 0.00 |
| Stomach | 0.03 ± 0.00 | 0.02 ± 0.01 | 0.03 ± 0.01 | 0.05 ± 0.03 | 0.03 ± 0.01 | 0.03 ± 0.01 | 0.02 ± 0.00 | 0.01 ± 0.00 |
| Liver | 0.43 ± 0.11 | 1.28 ± 0.05 | 1.65 ± 0.18 | 1.21 ± 0.07 | 0.58 ± 0.14 | 0.44 ± 0.10 | 0.30 ± 0.19 | 0.08 ± 0.02 |
| Spleen | 0.02 ± 0.02 | 0.02 ± 0.01 | 0.01 ± 0.00 | 0.02 ± 0.01 | 0.02 ± 0.01 | 0.01 ± 0.00 | 0.01 ± 0.01 | 0.01 ± 0.00 |
| Intestines | 0.03 ± 0.01 | 0.04 ± 0.01 | 0.05 ± 0.02 | 0.03 ± 0.01 | 0.03 ± 0.01 | 0.03 ± 0.00 | 0.03 ± 0.00 | 0.02 ± 0.00 |
| Kidneys | 0.83 ± 0.14 | 1.35 ± 0.23 | 1.59 ± 0.15 | 1.11 ± 0.14 | 0.84 ± 0.15 | 0.92 ± 0.06 | 0.33 ± 0.11 | 0.20 ± 0.07 |

**Table S2**. Decay-corrected % injected activities (%IA) per mass of tissue for 100µL of 740kBq [$^{111}$In]In-DTPA-dendrimers administered intravenously to BALB/c mice bearing **intracranial 4T1** triple negative breast tumors. Mice were **pretreated with cisplatin** (5 mg kg$^{-1}$, 24 h earlier). Error bars show the standard deviation for n = 2 mice per condition at each time point.

| % IA [$^{111}$In]In-DTPA-dendrimer | | | | | | | | |
|---|---|---|---|---|---|---|---|---|
| Time (hours) | 0.25 | 1 | 4 | 8 | 24 | 48 | 72 | 120 |
| Tumor | 0.06 ± 0.02 | 0.07 ± 0.04 | 0.08 ± 0.03 | 0.10 ± 0.02 | 0.14 ± 0.01 | 0.12 ± 0.01 | 0.06 ± 0.02 | 0.02 ± 0.01 |
| Brain | 0.04 ± 0.02 | 0.05 ± 0.01 | 0.04 ± 0.01 | 0.07 ± 0.01 | 0.05 ± 0.01 | 0.03 ± 0.01 | 0.03 ± 0.00 | 0.02 ± 0.01 |
| Blood | 41.63 ± 17.63 | 22.31 ± 3.83 | 19.37 ± 0.36 | 11.54 ± 4.94 | 2.15 ± 0.06 | 2.03 ± 0.04 | 1.60 ± 0.33 | 0.22 ± 0.03 |
| Heart | 1.58 ± 0.51 | 0.88 ± 0.27 | 0.24 ± 0.06 | 0.19 ± 0.00 | 0.12 ± 0.05 | 0.13 ± 0.08 | 0.10 ± 0.05 | 0.07 ± 0.04 |
| Lungs | 0.02 ± 0.01 | 0.02 ± 0.00 | 0.02 ± 0.00 | 0.02 ± 0.01 | 0.02 ± 0.01 | 0.01 ± 0.00 | 0.01 ± 0.00 | 0.01 ± 0.00 |
| Stomach | 0.03 ± 0.01 | 0.02 ± 0.00 | 0.02 ± 0.00 | 0.06 ± 0.02 | 0.04 ± 0.02 | 0.03 ± 0.01 | 0.03 ± 0.01 | 0.02 ± 0.00 |
| Liver | 0.54 ± 0.06 | 1.46 ± 0.16 | 1.86 ± 0.25 | 1.37 ± 0.12 | 0.72 ± 0.17 | 0.66 ± 0.29 | 0.54 ± 0.12 | 0.12 ± 0.02 |
| Spleen | 0.03 ± 0.01 | 0.02 ± 0.01 | 0.01 ± 0.01 | 0.02 ± 0.01 | 0.01 ± 0.00 | 0.01 ± 0.00 | 0.01 ± 0.00 | 0.02 ± 0.00 |
| Intestines | 0.04 ± 0.02 | 0.04 ± 0.02 | 0.06 ± 0.01 | 0.04 ± 0.00 | 0.03 ± 0.01 | 0.04 ± 0.01 | 0.04 ± 0.02 | 0.02 ± 0.01 |
| Kidneys | 0.99 ± 0.39 | 1.69 ± 0.40 | 1.79 ± 0.39 | 1.02 ± 0.35 | 0.93 ± 0.16 | 0.45 ± 0.07 | 0.39 ± 0.14 | 0.25 ± 0.17 |

**Table S3**. Decay-corrected % injected activities (%IA) per mass of tissue for 100µL of 740kBq [$^{111}$In]In-DTPA-dendrimers on BALB/c mice with **orthotopic (in the mammary fat pad) 4T1** Triple Negative Breast Cancer. Error bars correspond to standard deviations of n=2 mice per condition per time point.

| % IA [$^{111}$In]In-DTPA-dendrimer | | | | | | | | |
|---|---|---|---|---|---|---|---|---|
| Time (hours) | 0.25 | 1 | 4 | 8 | 24 | 48 | 72 | 120 |
| Tumor | 0.12 ± 0.02 | 0.61± 0.15 | 1.12± 0.03 | 1.93± 0.37 | 2.45± 0.21 | 2.29± 0.20 | 1.52± 0.25 | 0.79± 0.06 |
| Brain | 0.02 ± 0.00 | 0.03 ± 0.01 | 0.02 ± 0.00 | 0.03 ± 0.01 | 0.04 ± 0.00 | 0.02 ± 0.01 | 0.02 ± 0.00 | 0.01 ± 0.00 |
| Blood | 34.19 ± 3.03 | 17.11 ± 9.73 | 10.51 ± 4.18 | 5.45 ± 1.77 | 2.41 ± 0.05 | 1.32 ± 0.67 | 0.87 ± 0.19 | 0.16 ± 0.1 |
| Heart | 0.52 ± 0.19 | 0.29 ± 0.06 | 0.19 ± 0.00 | 0.13 ± 0.02 | 0.10 ± 0.01 | 0.09 ± 0.04 | 0.07 ± 0.01 | 0.05 ± 0.02 |
| Lungs | 0.02 ± 0.01 | 0.02 ± 0.00 | 0.02 ± 0.01 | 0.02 ± 0.00 | 0.03 ± 0.01 | 0.02 ± 0.01 | 0.02 ± 0.00 | 0.01 ± 0.01 |
| Stomach | 0.02 ± 0.01 | 0.03 ± 0.00 | 0.02 ± 0.00 | 0.02 ± 0.00 | 0.01 ± 0.00 | 0.01 ± 0.00 | 0.01 ± 0.00 | 0.01 ± 0.00 |
| Liver | 0.40 ± 0.19 | 0.72 ± 0.19 | 0.98 ± 0.04 | 0.50 ± 0.03 | 0.37 ± 0.09 | 0.17 ± 0.05 | 0.15 ± 0.06 | 0.08 ± 0.02 |
| Spleen | 0.01 ± 0.00 | 0.02 ± 0.00 | 0.03 ± 0.01 | 0.02 ± 0.00 | 0.02 ± 0.00 | 0.02 ± 0.00 | 0.01 ± 0.00 | 0.01 ± 0.00 |
| Intestines | 0.01± 0.01 | 0.04± 0.00 | 0.03± 0.00 | 0.02± 0.01 | 0.02± 0.01 | 0.01± 0.00 | 0.01± 0.00 | 0.01± 0.00 |
| Kidneys | 0.12 ± 0.00 | 0.37 ± 0.17 | 1.22 ± 0.45 | 1.41 ± 0.04 | 1.24 ± 0.09 | 0.52 ± 0.21 | 0.17 ± 0.08 | 0.11 ± 0.04 |

**Table S4**. Decay-corrected % injected activities (%IA) per mass of tissue for 100µL of 740kBq [$^{111}$In]In-DTPA-dendrimers administered intravenously to BALB/c mice bearing **orthotopic (in the mammary fat pad) 4T1** triple negative breast tumors. Mice were **pretreated with cisplatin** (5 mg kg$^{-1}$, 24 h earlier). Error bars show the standard deviation for n = 2 mice per condition at each time point.

| % IA [$^{111}$In]In-DTPA-dendrimer | | | | | | | | |
|---|---|---|---|---|---|---|---|---|
| Time (hours) | 0.25 | 1 | 4 | 8 | 24 | 48 | 72 | 120 |
| Tumor | 0.21 ± 0.04 | 1.11 ± 0.22 | 1.74 ± 0.27 | 2.51 ± 0.35 | 3.04 ± 0.30 | 2.87 ± 0.10 | 2.05 ± 0.17 | 1.20 ± 0.10 |
| Brain | 0.02 ± 0.00 | 0.03 ± 0.00 | 0.03 ± 0.01 | 0.03 ± 0.00 | 0.03 ± 0.00 | 0.03 ± 0.00 | 0.02 ± 0.00 | 0.02 ± 0.00 |
| Blood | 45.43 ± 10.50 | 20.54 ± 3.94 | 15.55 ± 1.69 | 7.66 ± 3.03 | 3.25 ± 1.29 | 1.59 ± 0.54 | 0.55 ± 0.03 | 0.18 ± 0.10 |
| Heart | 0.75 ± 0.05 | 0.47 ± 0.05 | 0.32 ± 0.10 | 0.16 ± 0.06 | 0.13 ± 0.02 | 0.10 ± 0.04 | 0.10 ± 0.02 | 0.10 ± 0.02 |
| Lungs | 0.02 ± 0.00 | 0.02 ± 0.00 | 0.03 ± 0.00 | 0.02 ± 0.00 | 0.02 ± 0.00 | 0.01 ± 0.00 | 0.01 ± 0.00 | 0.01 ± 0.00 |
| Stomach | 0.01 ± 0.00 | 0.02 ± 0.00 | 0.03 ± 0.00 | 0.02 ± 0.00 | 0.03 ± 0.00 | 0.02 ± 0.00 | 0.02 ± 0.00 | 0.01 ± 0.01 |
| Liver | 0.30 ± 0.09 | 0.84 ± 0.13 | 1.10 ± 0.35 | 0.74 ± 0.15 | 0.37 ± 0.15 | 0.17 ± 0.06 | 0.11 ± 0.02 | 0.09 ± 0.03 |
| Spleen | 0.02 ± 0.00 | 0.03 ± 0.00 | 0.03 ± 0.00 | 0.05 ± 0.00 | 0.03 ± 0.00 | 0.02 ± 0.00 | 0.01 ± 0.00 | 0.02 ± 0.00 |
| Intestines | 0.02 ± 0.00 | 0.02 ± 0.01 | 0.03 ± 0.00 | 0.03 ± 0.00 | 0.02 ± 0.00 | 0.02 ± 0.00 | 0.02 ± 0.01 | 0.01 ± 0.00 |
| Kidneys | 0.11 ± 0.01 | 0.32 ± 0.10 | 0.67 ± 0.18 | 1.66 ± 0.22 | 0.63 ± 0.11 | 0.37 ± 0.02 | 0.24 ± 0.13 | 0.14 ± 0.05 |

**Table S5**. Decay-corrected % injected activities (%IA) per mass of tissue for 100µL of 740kBq [$^{111}$In]In-DTPA-dendrimers on BALB/c mice with **subcutaneous 4T1** Triple Negative Breast Cancer. Error bars correspond to standard deviations of n=2 mice per condition per time point.

| % IA [$^{111}$In]In-DTPA-dendrimer | | | | | | | | |
|---|---|---|---|---|---|---|---|---|
| Time (hours) | 0.25 | 1 | 4 | 8 | 24 | 48 | 72 | 120 |
| Tumor | 0.16 ± 0.01 | 0.55 ± 0.14 | 0.96 ± 0.22 | 1.85 ± 0.41 | 3.06 ± 0.00 | 2.89 ± 0.17 | 2.56 ± 0.34 | 1.24 ± 0.02 |
| Brain | 0.01 ± 0.00 | 0.02 ± 0.01 | 0.02 ± 0.01 | 0.04 ± 0.01 | 0.02 ± 0.00 | 0.02 ± 0.00 | 0.02 ± 0.01 | 0.02 ± 0.01 |
| Blood | 31.67 ± 1.46 | 19.12 ± 4.72 | 4.76 ± 0.93 | 5.49 ± 1.25 | 2.32 ± 0.84 | 1.48 ± 0.32 | 0.60 ± 0.24 | 0.17 ± 0.02 |
| Heart | 0.60 ± 0.13 | 0.36 ± 0.08 | 0.27 ± 0.08 | 0.21 ± 0.03 | 0.12 ± 0.03 | 0.09 ± 0.01 | 0.09 ± 0.01 | 0.07 ± 0.01 |
| Lungs | 0.01 ± 0.00 | 0.01 ± 0.00 | 0.03 ± 0.00 | 0.03 ± 0.00 | 0.02 ± 0.00 | 0.02 ± 0.00 | 0.01 ± 0.00 | 0.01 ± 0.00 |
| Stomach | 0.02 ± 0.01 | 0.02 ± 0.01 | 0.03 ± 0.01 | 0.03 ± 0.01 | 0.03 ± 0.00 | 0.02 ± 0.01 | 0.01 ± 0.00 | 0.01 ± 0.00 |
| Liver | 0.34 ± 0.13 | 0.60 ± 0.03 | 1.61 ± 0.36 | 1.74 ± 0.27 | 0.68 ± 0.10 | 0.34 ± 0.04 | 0.19 ± 0.05 | 0.14 ± 0.03 |
| Spleen | 0.01 ± 0.00 | 0.01 ± 0.00 | 0.03 ± 0.01 | 0.02 ± 0.00 | 0.02 ± 0.01 | 0.02 ± 0.00 | 0.01 ± 0.00 | 0.02 ± 0.00 |
| Intestines | 0.01 ± 0.01 | 0.03 ± 0.02 | 0.03 ± 0.01 | 0.02 ± 0.01 | 0.02 ± 0.01 | 0.01 ± 0.00 | 0.01 ± 0.00 | 0.01 ± 0.01 |
| Kidneys | 0.31 ± 0.07 | 0.52 ± 0.09 | 0.97 ± 0.24 | 2.01 ± 0.02 | 1.44 ± 0.10 | 0.85 ± 0.18 | 0.30 ± 0.04 | 0.09 ± 0.04 |

**Table S6**. Decay-corrected % injected activities (%IA) per mass of tissue for 100µL of 740kBq [$^{111}$In]In-DTPA-dendrimers administered intravenously to BALB/c mice bearing **subcutaneous 4T1** triple negative breast tumors. Mice were **pretreated with cisplatin** (5 mg kg$^{-1}$, 24 h earlier). Error bars show the standard deviation for n = 2 mice per condition at each time point.

| % IA [$^{111}$In]In-DTPA-dendrimer | | | | | | | | |
|---|---|---|---|---|---|---|---|---|
| Time (hours) | 0.25 | 1 | 4 | 8 | 24 | 48 | 72 | 120 |
| Tumor | 0.22 ± 0.01 | 0.92 ± 0.11 | 1.18 ± 0.21 | 2.59 ± 0.41 | 3.69 ± 0.25 | 3.55 ± 0.21 | 3.30 ± 0.21 | 1.62 ± 0.13 |
| Brain | 0.01 ± 0.00 | 0.02 ± 0.01 | 0.02 ± 0.01 | 0.01 ± 0.00 | 0.03 ± 0.01 | 0.02 ± 0.00 | 0.01 ± 0.00 | 0.01 ± 0.00 |
| Blood | 37.00 ± 4.82 | 19.62 ± 3.04 | 11.56 ± 2.35 | 7.94 ± 1.50 | 2.67 ± 0.31 | 1.60 ± 0.09 | 1.01 ± 0.05 | 0.23 ± 0.04 |
| Heart | 0.69 ± 0.04 | 0.42 ± 0.03 | 0.24 ± 0.01 | 0.17 ± 0.03 | 0.11 ± 0.01 | 0.08 ± 0.03 | 0.07 ± 0.03 | 0.06 ± 0.00 |
| Lungs | 0.01 ± 0.00 | 0.01 ± 0.00 | 0.02 ± 0.00 | 0.03 ± 0.00 | 0.03 ± 0.00 | 0.02 ± 0.00 | 0.02 ± 0.00 | 0.01 ± 0.00 |
| Stomach | 0.01 ± 0.00 | 0.01 ± 0.00 | 0.02 ± 0.01 | 0.02 ± 0.01 | 0.03 ± 0.00 | 0.02 ± 0.01 | 0.01 ± 0.00 | 0.01 ± 0.00 |
| Liver | 0.46 ± 0.04 | 0.68 ± 0.00 | 1.38 ± 0.15 | 1.95 ± 0.53 | 1.65 ± 0.12 | 1.15 ± 0.29 | 0.44 ± 0.15 | 0.19 ± 0.07 |
| Spleen | 0.02 ± 0.00 | 0.02 ± 0.00 | 0.03 ± 0.01 | 0.03 ± 0.00 | 0.02 ± 0.00 | 0.01 ± 0.00 | 0.01 ± 0.00 | 0.01 ± 0.00 |
| Intestines | 0.01 ± 0.01 | 0.02 ± 0.00 | 0.02 ± 0.00 | 0.02 ± 0.00 | 0.01 ± 0.00 | 0.01 ± 0.00 | 0.01 ± 0.00 | 0.01 ± 0.00 |
| Kidneys | 0.42 ± 0.05 | 0.70 ± 0.02 | 1.50 ± 0.15 | 2.34 ± 0.09 | 1.33 ± 0.14 | 0.94 ± 0.15 | 0.47 ± 0.02 | 0.14 ± 0.04 |

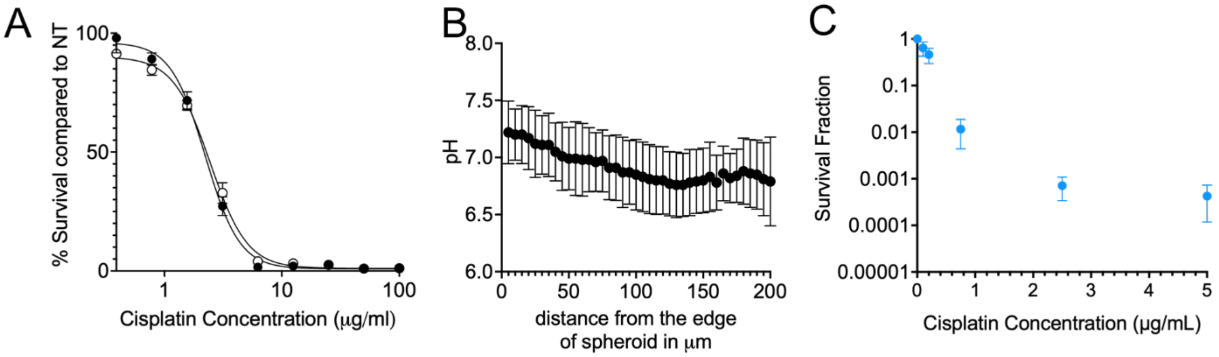


| Drug | pH | IC-50 |
|---|---|---|
| Cisplatin | 7.4 | 2.22 ± 0.07 µg/mL |
| Cisplatin | 6.5 | 2.45 ± 0.07 µg/mL |

**Fig. S1 (A)** Dose-response curves for 4T1 TNBC cells exposed to increasing cisplatin concentrations at pH 6.5 and pH 7.4. Error bars denote the standard deviation from three independent runs (n = 3). The $IC_{50}$ was evaluated at pH 6.5 representing the acidic pH in the spheroid core, as shown in Fig. S1 (B).

*Method*: *Cisplatin* $IC_{50}$

4T1 cells were seeded in 96-well plates for 12 hours prior to treatment, and were then incubated for 6 hours with cisplatin, at serial dilutions. Following treatment, the wells were washed twice with PBS and fresh RPMI 1640 media supplemented with 10% FBS and 1% PS was added. The cells were then allowed to grow for 28 hours (equivalent to two doubling times). Cell viability was assessed using the 3-(4,5-dimethylthiazol-2-yl)-2,5-diphenyltetrazolium bromide (MTT) assay according to the manufacturer's instructions, with absorbance measured at 570 nm using a Spectramax M3 Microplate Reader (Molecular Devices, San Jose, CA).

**Fig. S1 (B)** Measurement of 4T1 spheroids' interstitial pH by employing the membrane impermeant, pH indicator SNARF as previously described [1].

**Fig. S1 (C)** clonogenic survival of 4T1 cells exposed to cisplatin. Method as described in main text.

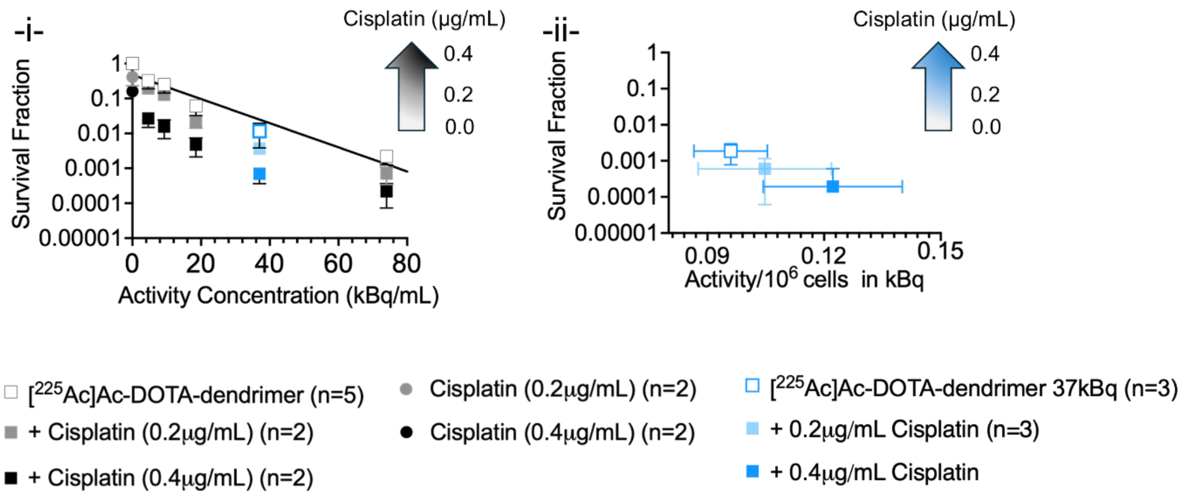


**Fig. S2 (A)** The clonogenic survival of 4T1 cells exposed to **[$^{225}$Ac]Ac-DOTA-dendrimers** decreased in the presence of cisplatin independent of the activity concentrations during incubation (-i-), and, lower extents of clonogenic survival correlated with higher levels of activity associated per cell, that increased with increasing concentrations of cisplatin at same activity concentrations during incubation (-ii-). Clonogenic survival of 4T1 triple-negative breast cancer cells following a 6 h incubation at 37 °C with increasing activities of [$^{225}$Ac]Ac-DOTA-dendrimers, with or without cisplatin is shown in (-i-). (-ii-) Survival at the 37kBq/mL (1 µCi/mL) activity (indicated in panel -i- by the symbols in blue) replotted versus radioactivity per $10^6$ cells in the presence and absence of cisplatin. The dendrimer mass concentration was kept constant at 10µg/mL. Data represent mean ± standard deviation from n = 3 independent experiments.

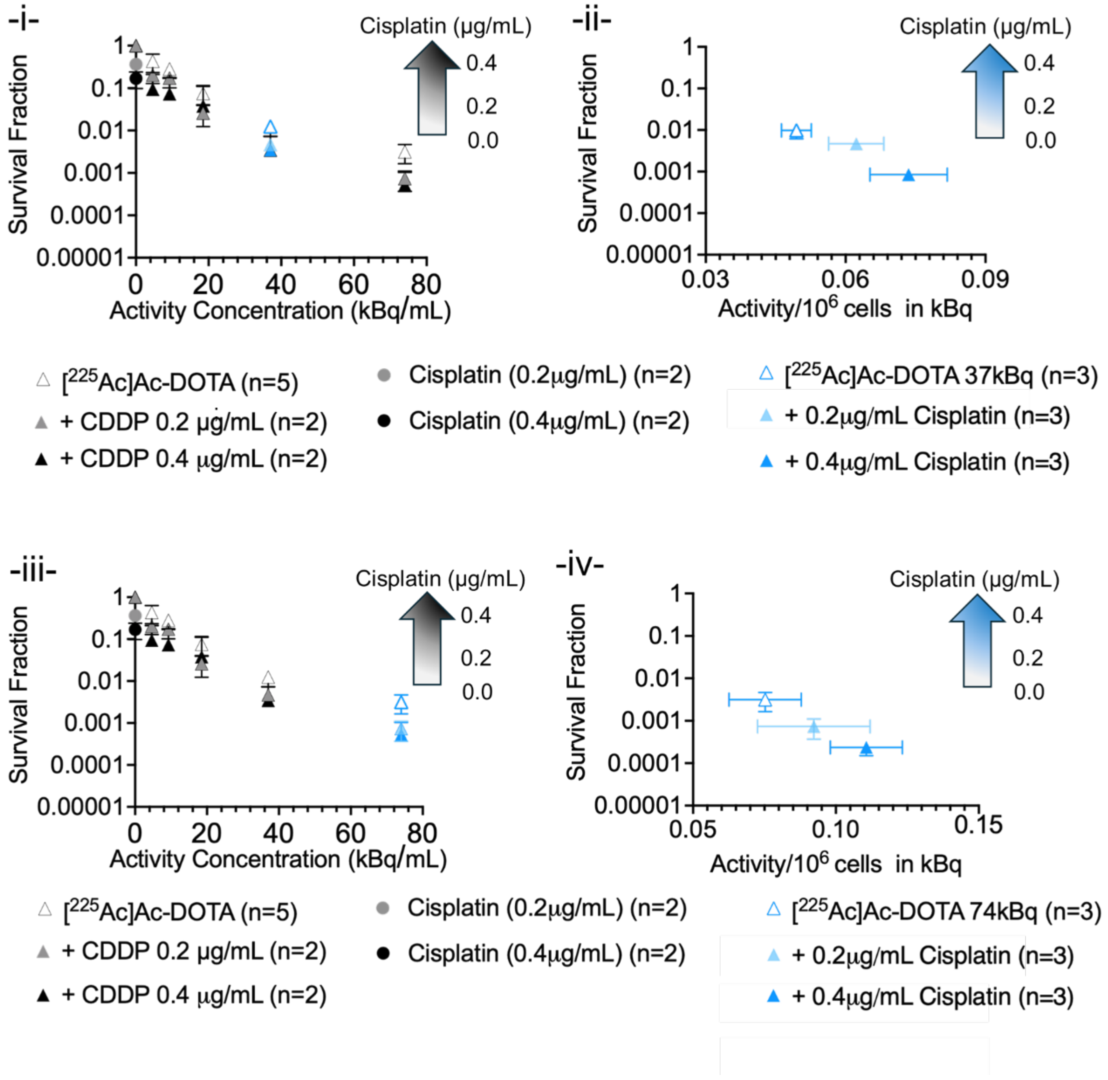


**Fig. S2 (B)** Clonogenic survival of 4T1 cells exposed to **[$^{225}$Ac]Ac-DOTA** (-i- and -iii- are the same plot indicating a different set of data re-plotted in -ii- and -iv-, respectively) and correlation of survival to activity associated per cell for different cisplatin concentrations (-ii-, -iv-). Clonogenic survival of 4T1 triple-negative breast cancer cells following a 6 h incubation at 37 °C with increasing activities of [$^{225}$Ac]Ac-DOTA, with or without cisplatin is shown in -i- and -iii-. Plots in -ii- and -iv- demonstrate the survival at the 37kBq/mL (1 µCi/mL) and 74kBq/mL (2 µCi/mL) activity (indicated in panel -i- and -ii-, respectively, by the symbols in blue) replotted versus activity per $10^6$ cells in the presence and absence of cisplatin. The dendrimer mass concentration was kept constant at 10 µg/mL. Data represent mean ± standard deviation from n = 3 independent experiments.

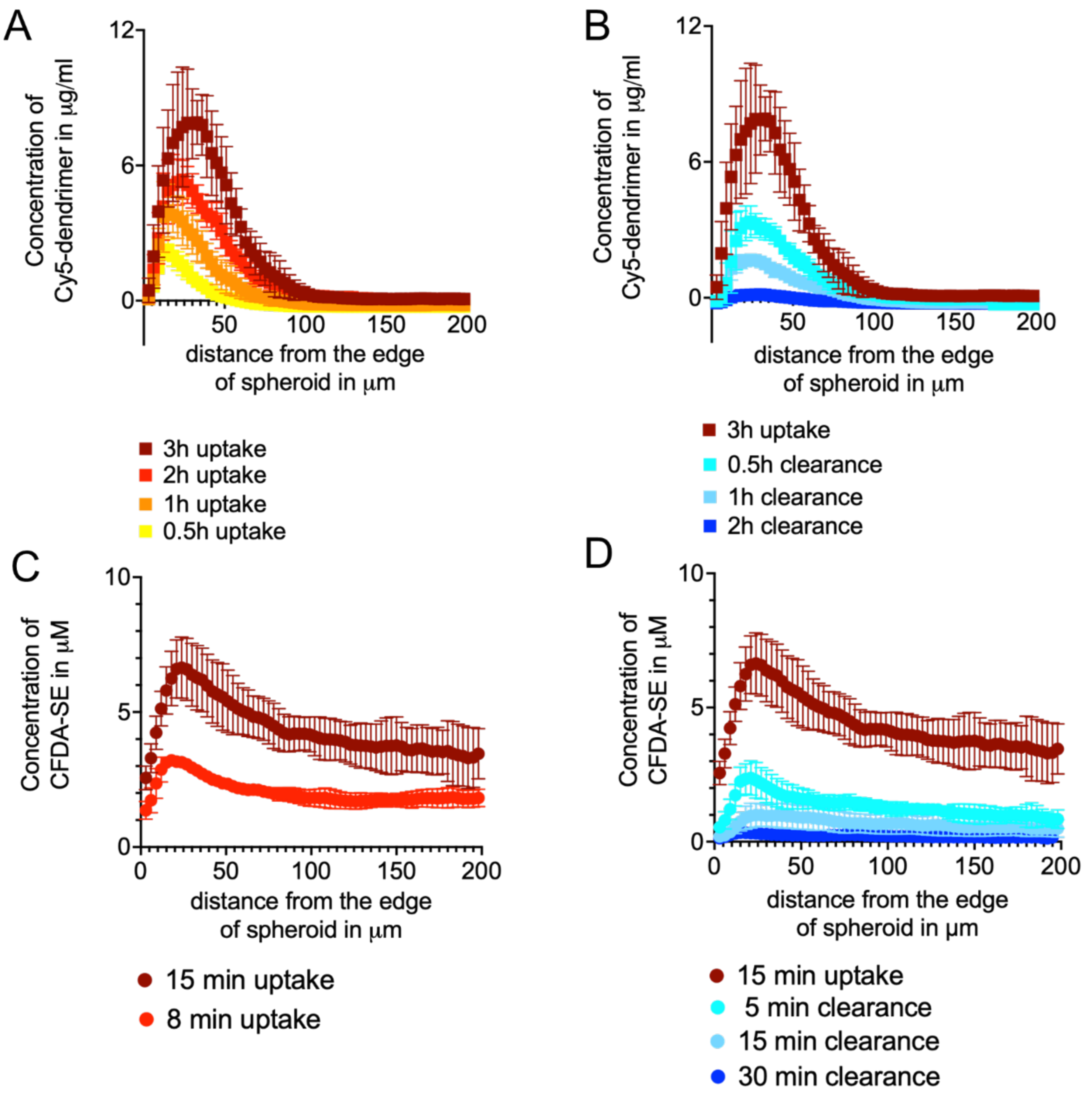


**FIG. S3.** Spatiotemporal profiles of Cy5-dendrimer (A, B) and CFDA-SE (C, D), employed as fluorescent surrogate of Cisplatin, in 4T1 spheroids of 400μm diameter. Data points indicate the mean values ± the standard deviations of n=3 different spheroids per time point.

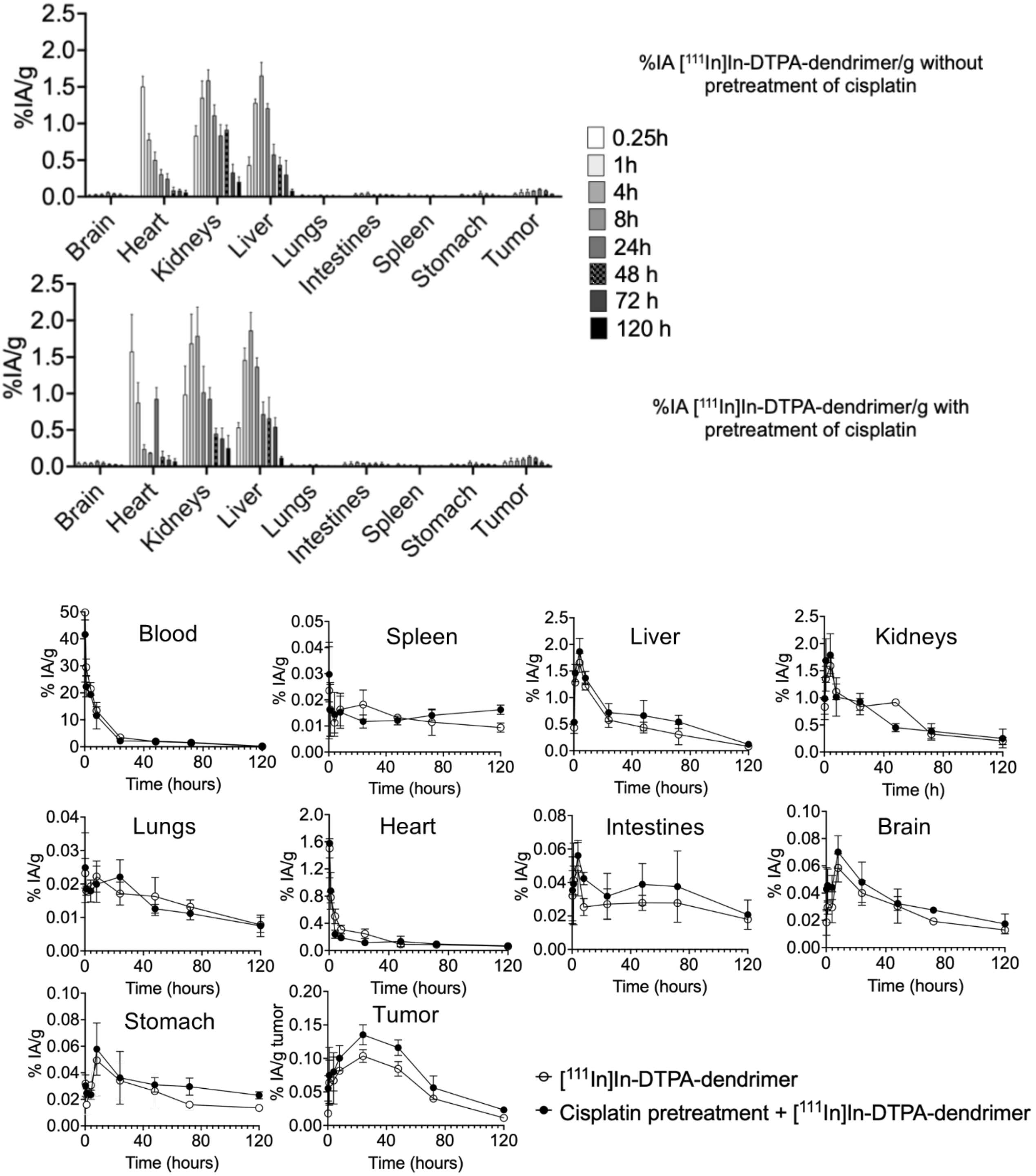


**Fig. S4 (A)** Biodistributions of systemically administered [$^{111}$In]In-DTPA-dendrimers in Balb/c mice bearing **intracranial 4T1 TNBC tumors** without cisplatin pretreatment (top panel or white symbols) and after intraperitoneal administration of cisplatin (5 mg/Kg) injected 24 hours earlier (bottom panel or black symbols). Values are mean ± SD for n = 2 mice per time point.

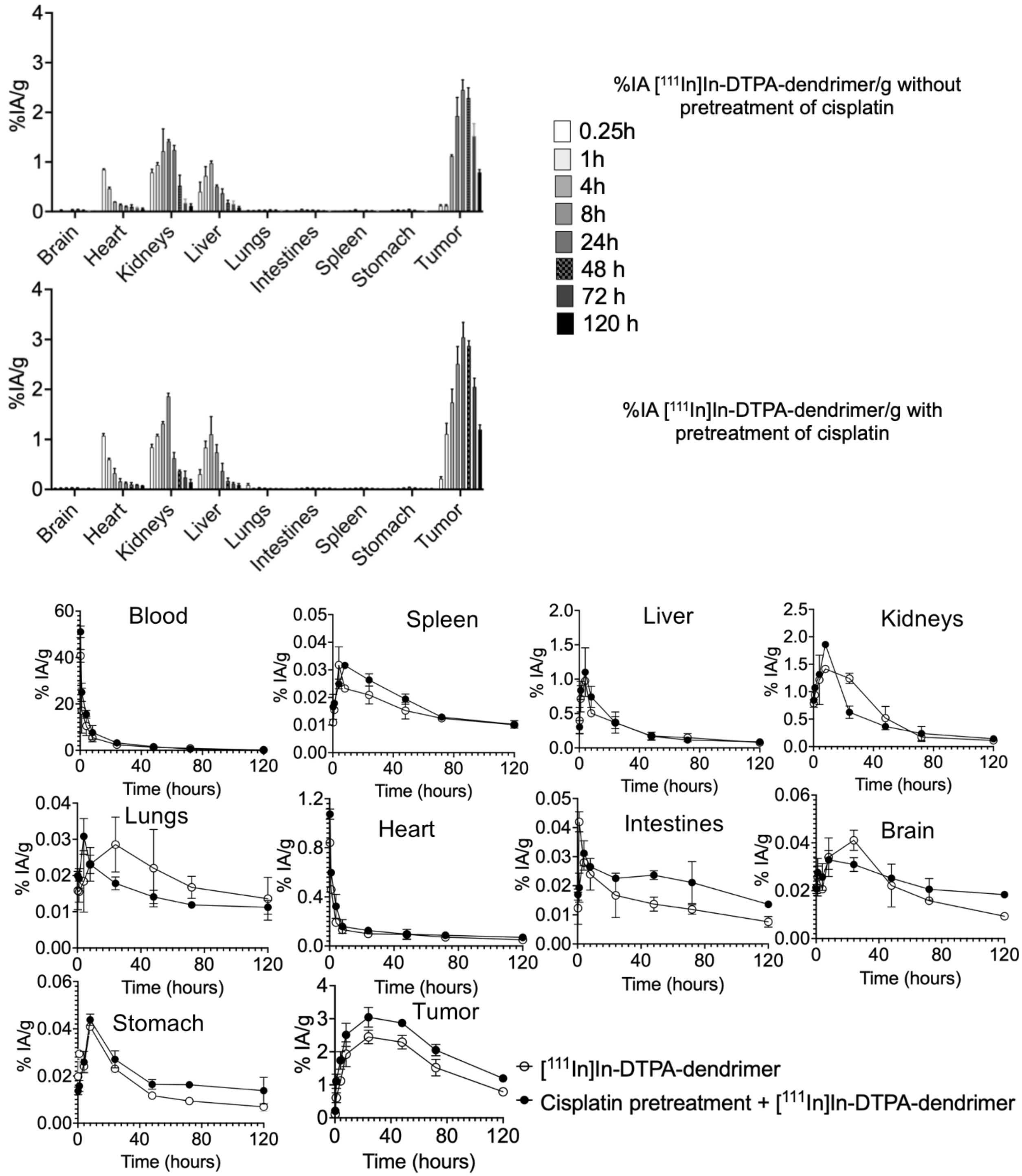


**Fig. S4 (B)** Biodistributions of systemically administered [$^{111}$In]In-DTPA-dendrimers in Balb/c mice bearing **4T1 TNBC tumors in the mammary fat pad** without cisplatin pretreatment (top panel or white symbols) and after intraperitoneal administration of cisplatin (5 mg/Kg) injected 24 hours earlier (bottom panel or black symbols). Values are mean ± SD for n = 2 mice per time point.

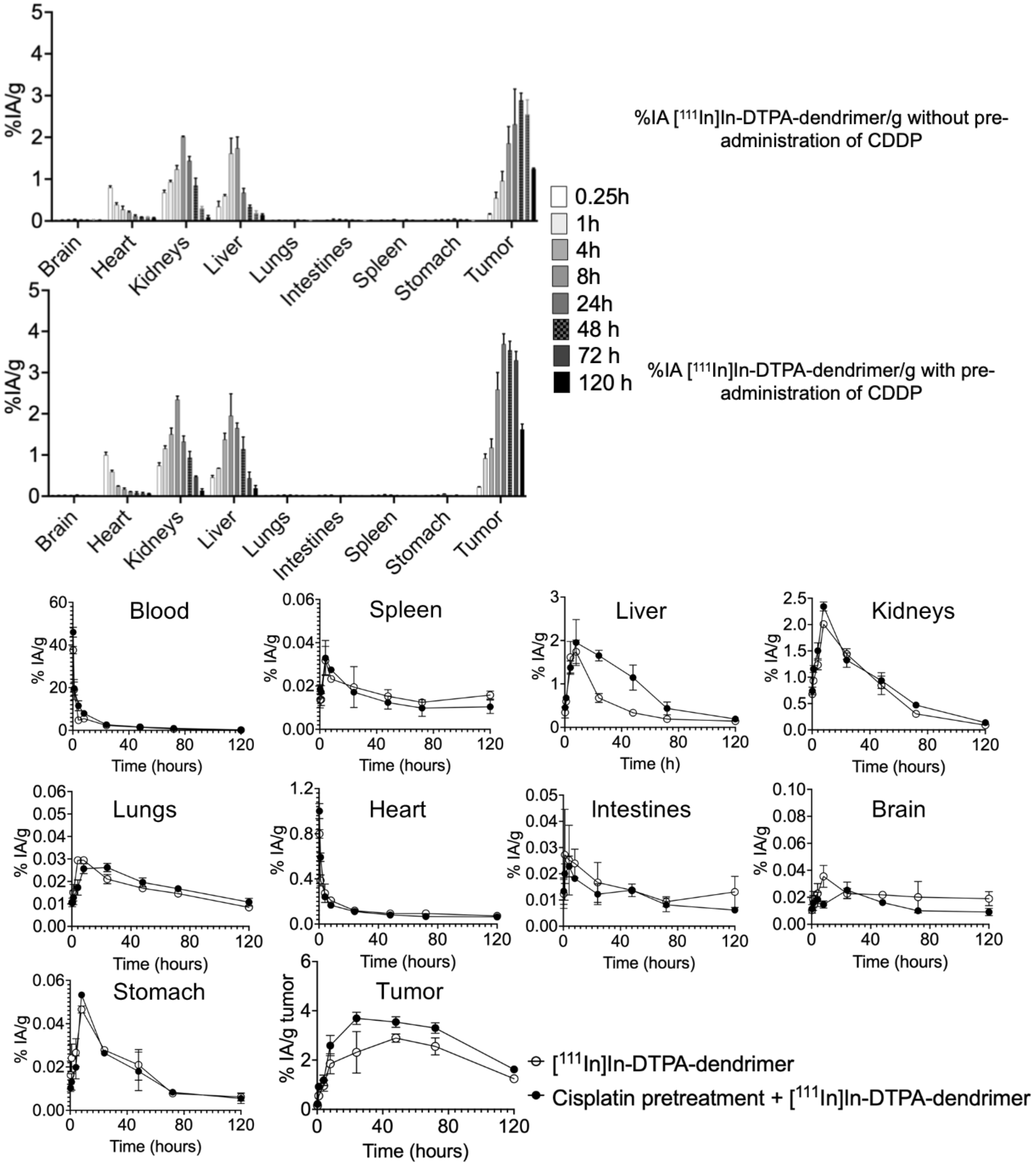


**Fig. S4 (C)** Biodistributions of systemically administered [$^{111}$In]In-DTPA-dendrimers in Balb/c mice bearing **subcutaneous 4T1 TNBC tumors** without cisplatin pretreatment (top panel or white symbols) and after intraperitoneal administration of cisplatin (5 mg/Kg) injected 24 hours earlier (bottom panel or black symbols). Values are mean ± SD for n = 2 mice per time point.

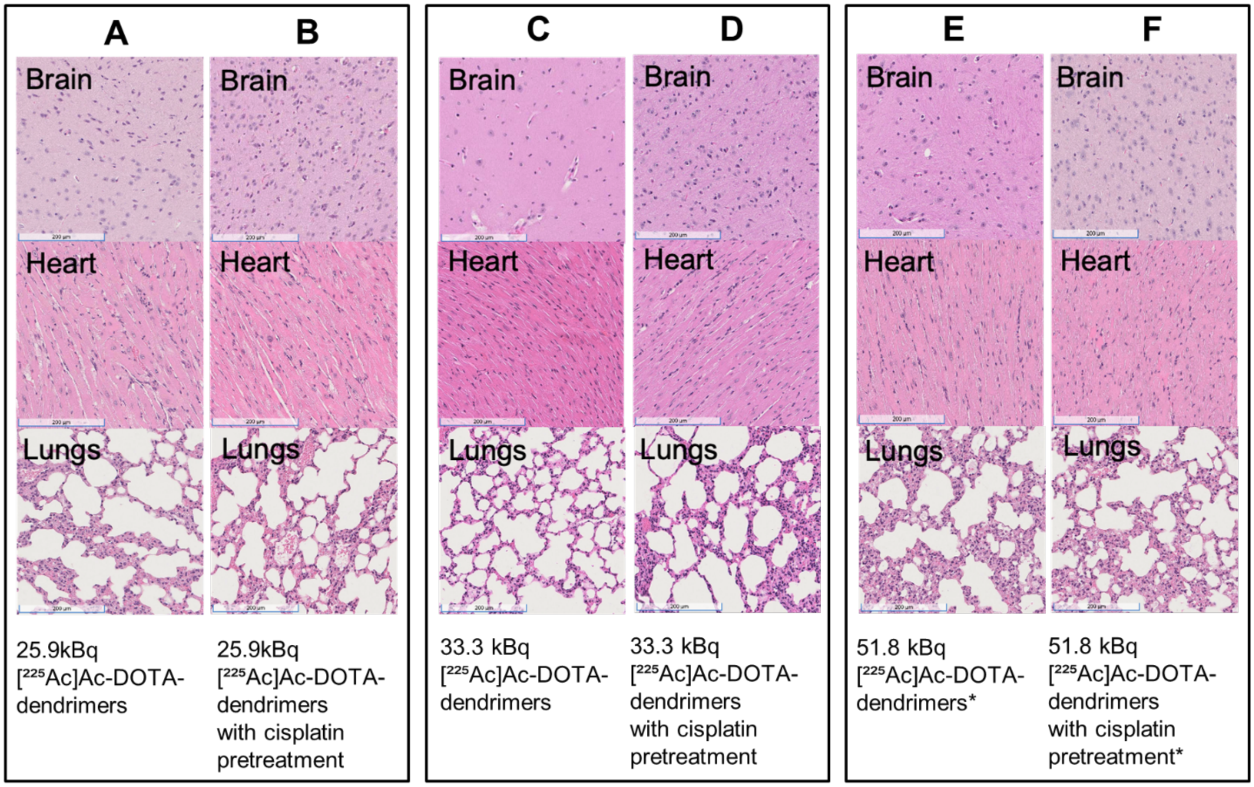


**Fig. S5** Acute toxicities in Balb/c tumor-free mice were not observed at any of the systemically injected activities of [$^{225}$Ac]Ac-dendrimers (with and without pretreatment with low-dose cisplatin). The onset of long-term toxicities (10-months after injection) was observed by histopathology, at systemically injected activities of 51.8 kBq. The mice didn't show any physical signs of toxicity. Six-week-old mice were treated with 25.9, 33.3 and/or 51.8kBq [$^{225}$Ac]Ac-DOTA-dendrimers/18g mouse, with or without cisplatin pretreatment (5mg/Kg) administered 24 hours earlier. During the 10-month observation period, none of the treated mice reached any of the study's endpoints. Ten months after administration of activity, mice were euthanized, and organs were harvested for histopathological analysis using H&E staining. The scale bar corresponds to 200µm on all images.

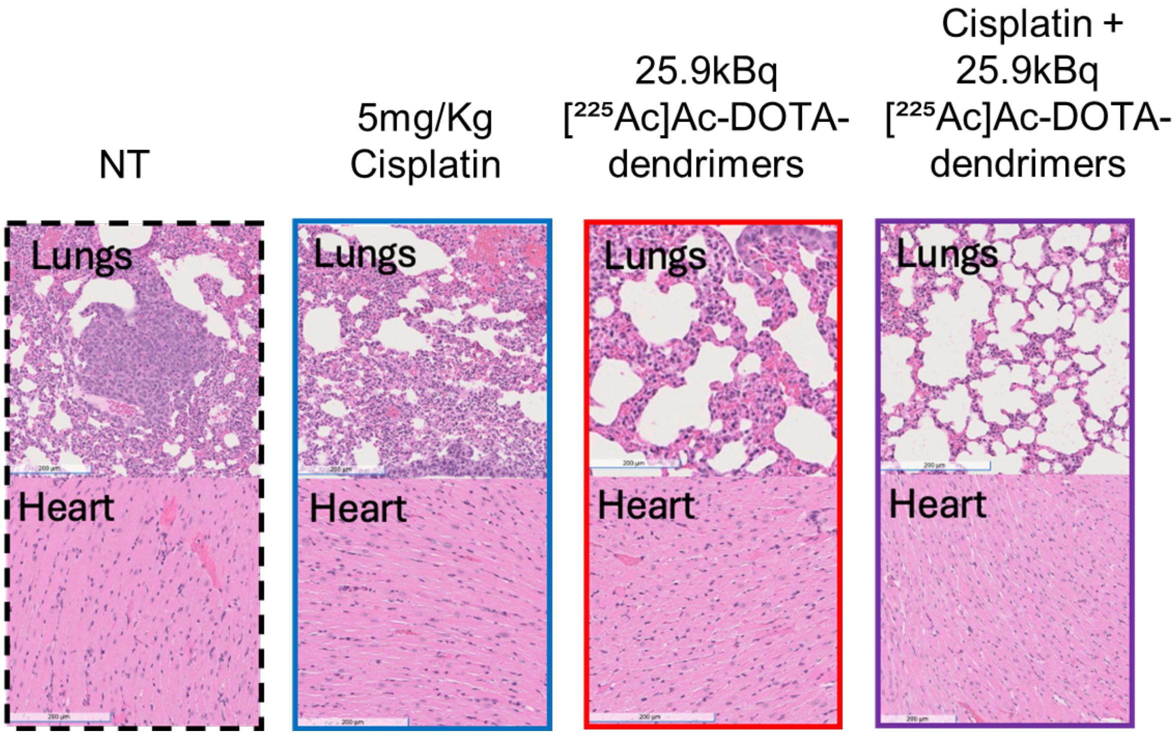


**Fig. S6** Balb/c mice with syngeneic 4T1 triple negative breast cancer implanted intracranially (from the study shown in Fig. 6): H&E-stained sections of lungs and hearts of mice sacrificed after indicated treatment. No significant toxicity was observed. Scale bar=200µm.

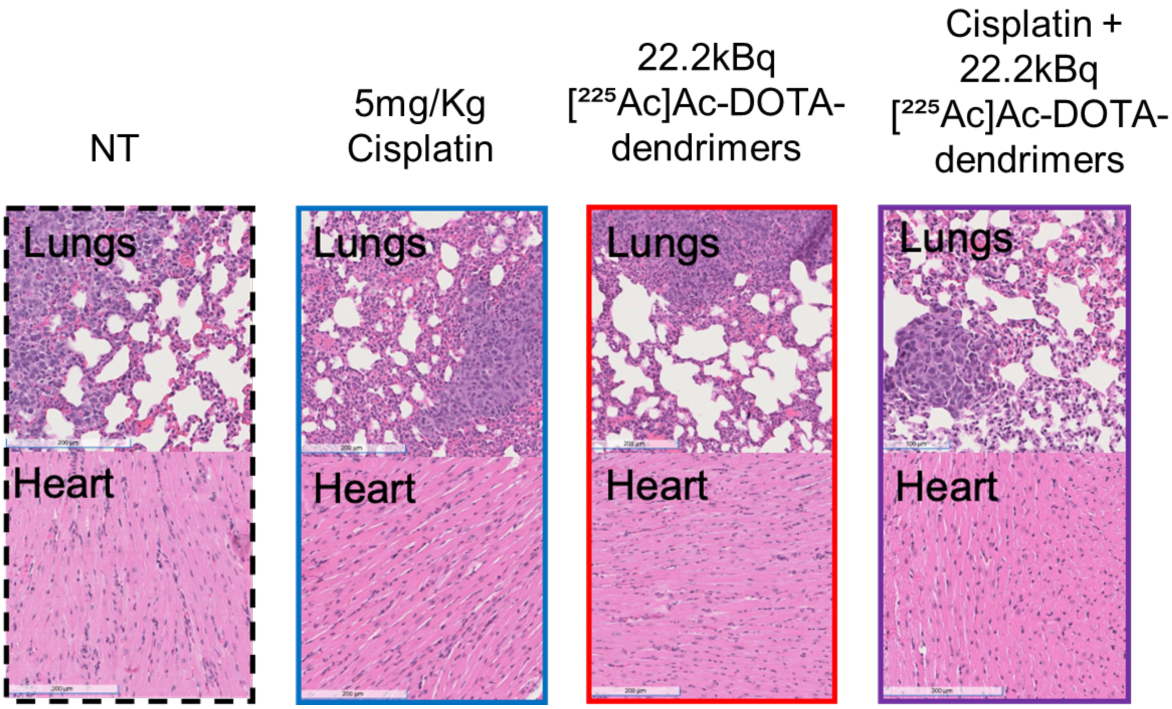


**Fig. S7** Balb/c mice with syngeneic 4T1 triple negative breast cancer implanted in the mammary fat pad (from the study shown in Fig. 7): H&E-stained sections of lungs and hearts of mice sacrificed after indicated treatment. No significant toxicity was observed. Scale bar=200µm.

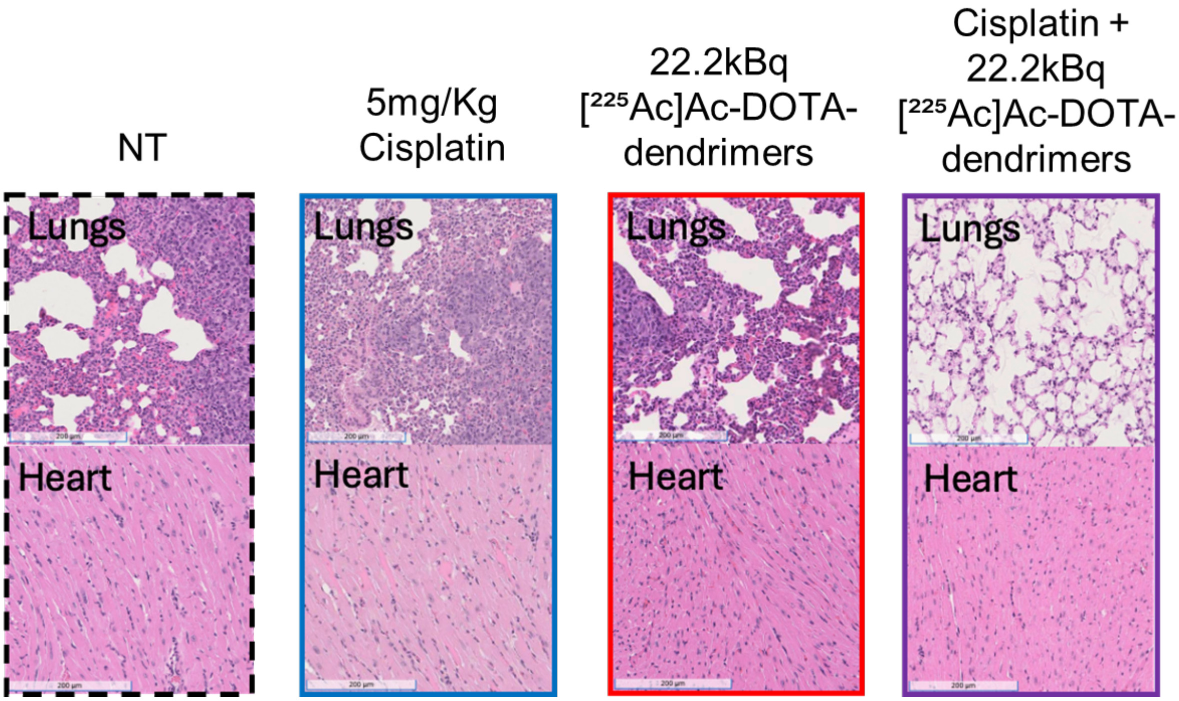


**Fig. S8** Balb/c mice with syngeneic 4T1 triple negative breast cancer implanted subcutaneously (from the study shown in Fig. 8): H&E-stained sections of lungs and hearts of mice sacrificed after indicated treatment. No significant toxicity was observed. Scale bar=200µm.

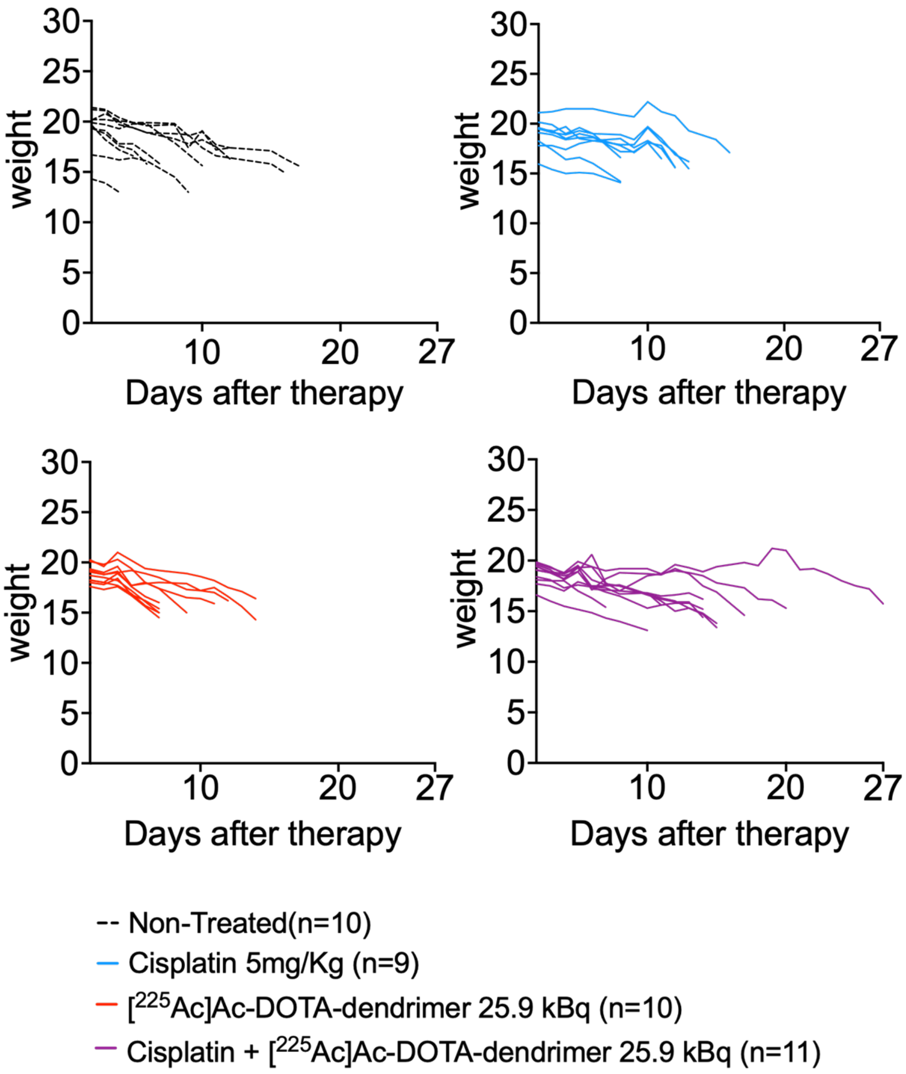


**Fig. S9** Weight of individual animals in the different treatment cohorts that were monitored as part of the tumor growth control studies in the **intracranial tumor model.** Animals were euthanized when weight loss was ≥ 20%.

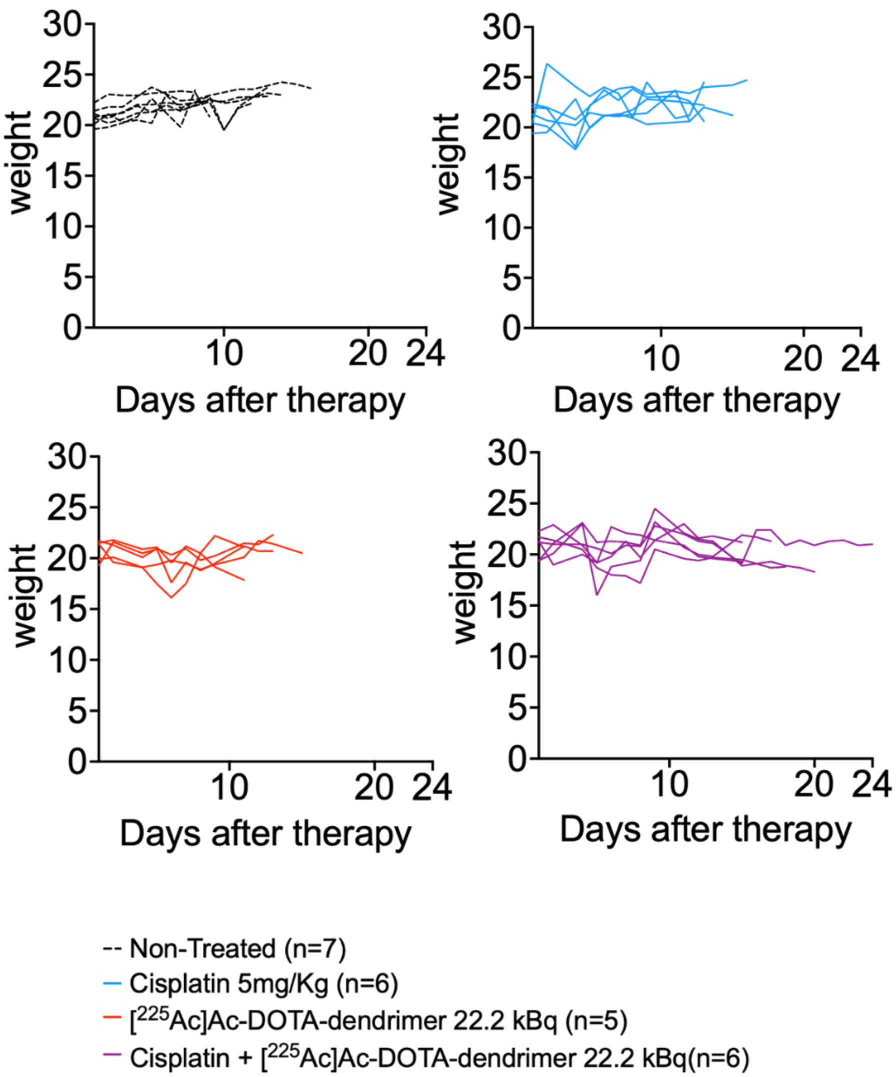


**Fig. S10** Weight of individual animals in the different treatment cohorts that were monitored as part of the 4T1 tumor growth control studies in the **mammary fat pad tumor model**. Animals were euthanized if tumor size was greater than 10% of body weight at the day of injection of therapy

Mass of Tumor (mg) = Tumor Volume ($mm^3$), and percentage was calculated as $\frac{100\times (Tumor\ mass\ (weight))}{(Combined\ body\ weight\ with\ tumor - tumor\ weight)_{day\ of\ injection}}$, or when the tumor interfered with the movement or function of mice or when weight loss was $\geq$ 20%.

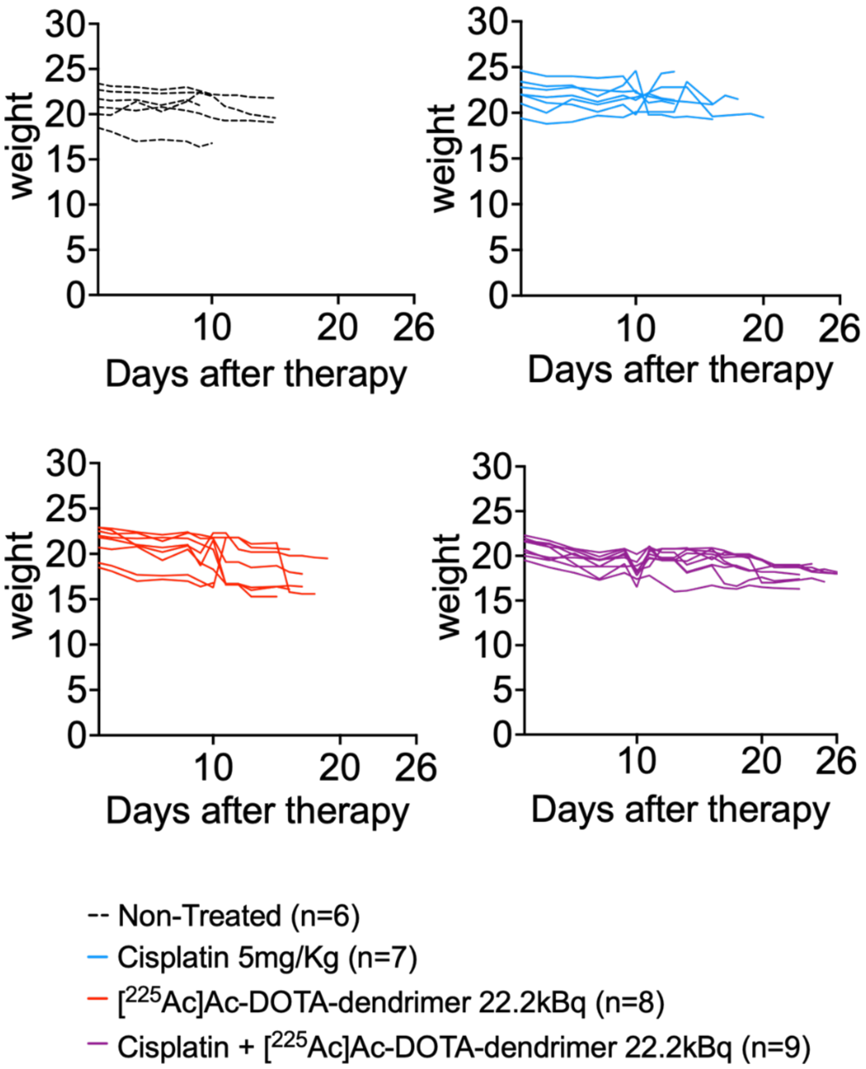


**Fig. S11** Weight of individual animals in the different treatment cohorts that were monitored as part of the 4T1 tumor growth control studies in the **subcutaneous tumor model**. Mice were euthanized, when the tumor interfered with the mobility and functioning of mice or when weight loss was ≥ 20%.
